# Significant variations in tolerance to clothianidin and pirimiphos-methyl in *Anopheles gambiae* and *Anopheles funestus* populations during a dramatic malaria resurgence despite sustained indoor residual spraying in Uganda

**DOI:** 10.1101/2025.02.13.638152

**Authors:** Ambrose Oruni, Emmanuel Arinaitwe, James Adiga, Geoffrey Otto, Patrick Kyagamba, Joseph Okoth, Daniel Ayo, Jackson Rwatooro Asiimwe, Maato Zedi, John Rek, Kyle J Walker, Ashlee Braithwaite, Jonathan Kayondo, Melissa D Conrad, Teun Bousema, Mark J I Paine, Hanafy M Ismail, Paul Krezanoski, Charles S. Wondji, Moses R. Kamya, Grant Dorsey, Martin J. Donnelly

## Abstract

A dramatic malaria resurgence occurred in areas of Uganda between 2020 and 2022 coincident with the switch to clothianidin-based formulations for indoor residual spraying. During the resurgence, *Anopheles funestus* numbers increased but when an alternative insecticide, pirimiphos methyl, was reintroduced in 2023, both malaria cases and *An. funestus* mosquito density fell. In this study, we investigated possible causes of the resurgence by assessing; 1) whether sufficient quantities of insecticide were sprayed; 2) the residual insecticide bioefficacy against wild mosquitoes and; 3) the insecticide susceptibility of vector populations using standard test tube assays and wall cone assays. In 2023, after adjusting for extraction efficiency, 70-80% of the houses had optimal residual concentrations of insecticides (clothianidin >0.3g/m^2^; pirimiphos methyl >0.5g/m^2^) with significant variations between sampling rounds and wall types. Mud walls had the lowest residual concentration of insecticides, and the lowest observed mortality in wall cone assays, compared to fired bricks with plaster/cement/paint. In the studies of residual bio efficacy, by World Health Organization (WHO) definitions, *An. funestus* showed resistance to clothianidin (<80% mortality) up to 11 months and susceptibility to pirimiphos methyl (>90% mortality) when exposed to wall surfaces up to 7 months post-spray. In WHO tube tests, variations were observed in susceptibility to clothianidin in *An. funestus* populations using dose- and time-response assays (80-98% mortality). In 2022, *An. gambiae* was largely susceptible to the clothianidin-based formulation Sumishield (85-90% mortality) although the levels dropped slightly in 2023 (60-85% mortality) mainly in mud and pole houses. In contrast, *An. gambiae* was mildly susceptible to the pirimiphos methyl-based formulation Actellic (∼80% mortality) and time response assays showed *An. gambiae populations* had very low knockdown and mortality at lower exposure time compared to *An. funestus*. Regression models showed a positive association between residual insecticide concentration (RIC) and mortality in houses sprayed with Sumishield but not Actellic houses.

Despite the possible variations observed in spray operations, the study revealed that *An*. *funestus* exhibited a higher tolerance to clothianidin-based formulations compared to *An. gambiae*, and this might have driven the malaria resurgence observed in Uganda. However, there are signals of *An. gambiae* resistance to pirimiphos-methyl which will require further investigation and monitoring.

## Introduction

Malaria control and elimination campaigns have largely relied on vector control interventions, especially in high transmission areas, mainly through long-lasting insecticidal nets (LLINs) and indoor residual spraying (IRS) (WHO, 2019). Historically, IRS played a vital role in campaigns aimed at eliminating malaria but its use in sub-Saharan Africa has been limited due to scarce resources. Until recently, IRS has relied on a limited range of insecticides namely; organochlorines, pyrethroids, carbamates and organophosphates. More recently, the World Health Organization (WHO) approved new classes of insecticides such as neonicotinoids (e.g. clothianidin) and pyrroles (e.g. chlorfenapyr) (WHO, 2022a) which has provided a needed expansion of the armamentarium. In Uganda, IRS has focused on high transmission zones since 2009, using a diverse array of formulations starting with carbamates (Bendiocarb) and then moving to the organophosphate ActellicⓇ 300 CS (referred here in as Actellic) – containing pirimiphos-methyl (Kamya et al., 2024). In 2020-2022, Uganda switched to clothianidin-based formulations; FludoraⓇ Fusion (referred here in as Fludora Fusion) and SumishieldⓇ 50 WG (referred here in as Sumishield). Clothianidin is a neonicotinoid with low mammalian toxicity, primarily used against insects of agricultural and veterinary importance (Simon-Delso et al., 2015). The mode of action of neonicotinoids is to target the nicotinic acetylcholine receptor (nAChR) in the insect central nervous system (Tomizawa & Casida, 2005). Sumishield has only clothianidin as the active ingredient while Fludora Fusion combines clothianidin with a pyrethroid – deltamethrin. An experimental hut trial in Benin showed that clothianidin in combination with deltamethrin induced high, long-lasting mosquito mortality (Ngufor et al., 2017), although the impact on clinical malaria indicators was not measured. In a survey of 16 countries, including Uganda, before the widespread use of clothianidin-based formulations for IRS, *Anopheles* vectors were largely susceptible to clothianidin (Oxborough et al., 2019). However, clothianidin resistance has recently been reported in Cameroon (2019-20) in *An. gambiae s.s* populations (Fouet et al., 2024) which contrasted with the susceptibility largely observed in *An. funestus* (Assatse et al., 2023). Therefore, it was predicted that clothianidin-based formulations would be effective for IRS in Uganda. However, a resurgence of malaria burden occurred following a change to clothianidin-based IRS (cIRS) in 5 districts of Uganda (Epstein, Maiteki-Ssebuguzi, et al., 2022). Our team continued to monitor this trend in a cohort of household from one of these districts (Tororo) where malaria incidence increased more than 8-fold (0.36 vs. 2.97 episodes per person-year, *P* < 0.0001) from 2021 to 2022 concomitantly with the emergence of *An. funestus* as the dominant vector (Kamya et al., 2024). Furthermore, when the programme switched back to pirimiphos methyl IRS (pIRS) in 2023, there was a marked decrease in malaria incidence coinciding with a decrease in *An. funestus* vector density (Kamya et al., 2024). For this study, we conducted a two-year longitudinal survey (2022-2023) to investigate potential causes of the resurgence with the hypotheses that the increase in malaria cases was at least partially due to; 1) sub-optimal insecticide deployment and/or 2) emergent physiological or behavioral resistance to clothianidin in the primary malaria vectors.

## Methods

### Ethical statement

Ethical approval was obtained from the Makerere University School of Medicine Research and Ethics Committee (REF 2019–134), the Uganda National Council for Science and Technology (HS 2700), the London School of Hygiene & Tropical Medicine Ethics Committee (17777), and the University of California, San Francisco Committee on Human Research (257790). Written informed consent was obtained for all participants before enrolment into the parent study.

### Study setting, characteristics of households and vector control interventions

The study was conducted in Tororo district where IRS has been conducted at least annually since 2014 and in Busia, a contiguous district, where IRS has never been deployed (Figure 1A). A total of 60 households from Tororo and 15 households from Busia (Figure 1B) were included in this study and enrolled in “The PRISM Border Cohort” parent study (Kamya et al., 2024). Household characteristics were recorded including the type of walls of the house (Table 1). The history of IRS campaigns has been fully described (Kamya et al., 2024). In this study, we focus on the period 2020-2023. Briefly, Fludora Fusion (clothianidin plus deltamethrin) was sprayed in March 2021, followed by Sumishield (clothianidin alone) in March 2022 and then a reversion to primarily Actellic in March 2023 with a few houses receiving Sumishield. In our study, 51 out of 60 houses were sprayed with Sumishield in 2022 while in 2023, 9 out of 60 houses were sprayed with Sumishield and 49 out of 60 houses were sprayed with Actellic (Table 1, Figure 1B). Whilst we followed the same sixty households throughout the study, home improvements resulted in some households moving between categories in Table 1.

**Figure. 1:**
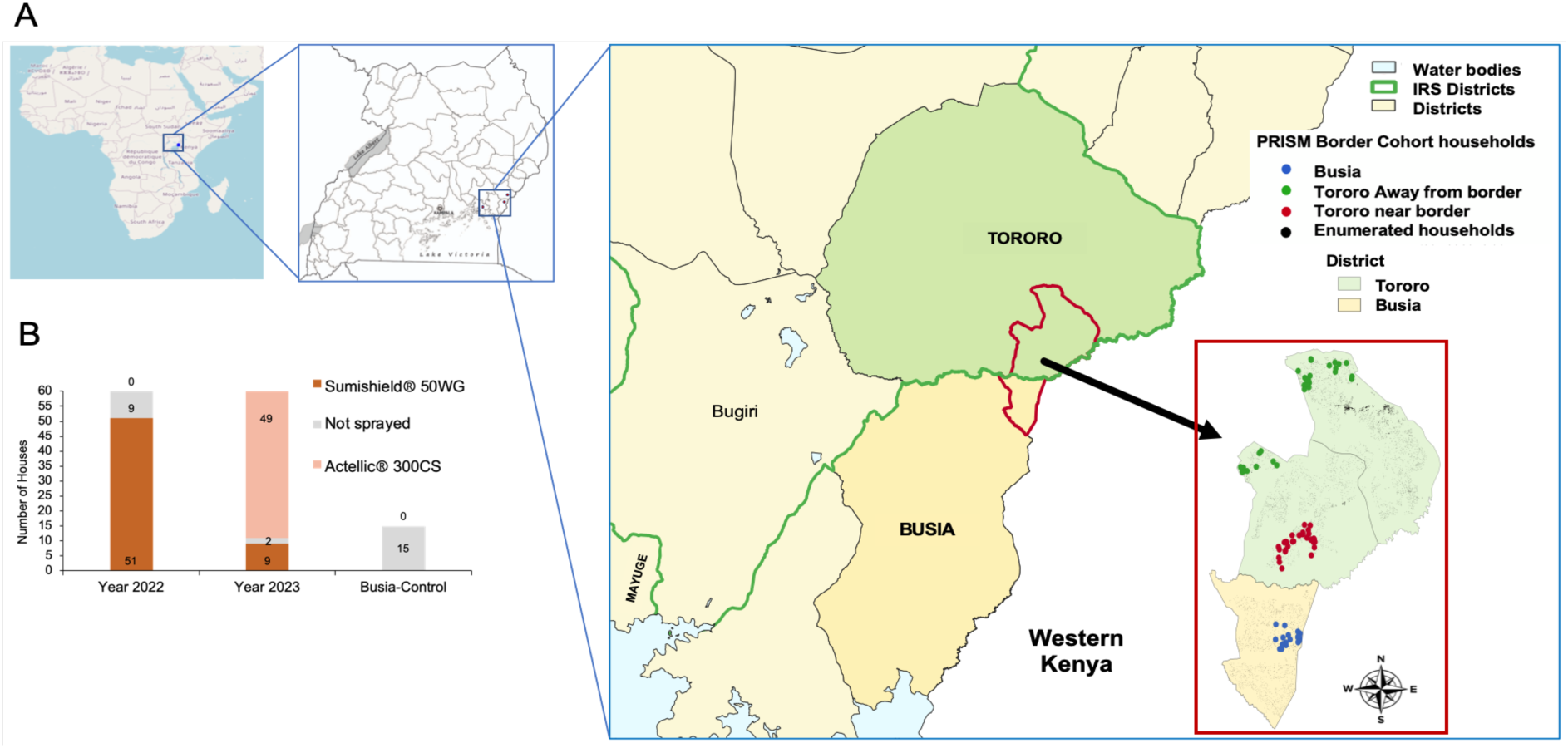
Location and number of households enrolled in the study. Map showing the location of study sites in Tororo and Busia districts (A) and distribution of the number of houses sprayed with either Sumishield or Actellic in 2022 and 2023 together with control houses from Busia district included in this study (B).

**Table 1:** Summary of house characteristics from the total number of households enrolled in the study from Tororo district.

| No. | Wall type | Total | Sprayed | Actellic® 300CS | Sumishield® 50WG | Unsprayed |
| --- | --- | --- | --- | --- | --- | --- |
| **<u>2022 IRS Tororo</u>** |  |  |  |  |  |  |
| 1. | Mud and poles | 34 | 31 | 0 | 31 | 3 |
| 2. | Burnt bricks with plaster-cement | 23 | 17 | 0 | 17 | 6 |
| 3. | Burnt bricks with mud | 3 | 3 | 0 | 3 | 0 |
| 4. | Burnt bricks with plaster-only | 0 | 0 | 0 | 0 | 0 |
| 5. | Burnt bricks with plaster-painted | 0 | 0 | 0 | 0 | 0 |
| **Total number of houses** |  | **60** | **51** | **0** | **51** | **9** |
| **<u>2022/23 Busia (Non-IRS)</u>** |  |  |  |  |  |  |
| 1. | Bricks cement-plaster | 6 | 0 | 0 | 0 | 6 |
| 2. | Mud or unburnt bricks | 9 | 0 | 0 | 0 | 9 |
| **Total number of houses** |  | **15** | **0** | **0** | **0** | **15** |
| **<u>2023 IRS Tororo</u>** |  |  |  |  |  |  |
| 1. | Mud and poles | 32 | 32 | 24 | 8 | 0 |
| 2. | Burnt bricks with plaster-cement | 22 | 21 | 20 | 1 | 1 |
| 3. | Burnt bricks with mud | 3 | 3 | 3 | 0 | 0 |
| 4. | Burnt bricks with plaster-only | 2 | 1 | 1 | 0 | 1 |
| 5. | Burnt bricks with plaster-painted | 1 | 1 | 1 | 0 | 0 |
| **Total number of houses** |  | **60** | **58** | **49** | **9** | **2** |

### Mosquito strains, collections and rearing

#### An. gambiae mosquitoes

In 2022, we used a combination of field mosquitoes raised from collected larvae (F0), the BusiaUg colony (*An. gambiae s.s*) that had been maintained in Nagongera insectary without selection, and the susceptible Kisumu lab strain (*An. gambiae s.s*) as a control. For mosquitoes collected from the field, we prioritised the use of adult females derived from larvae collected from breeding sites surrounding houses in areas where IRS was conducted. Molecular species identification on a sample of the adults raised revealed that most were *An. arabiensis* (55.4%) or *An. gambiae s.s.* (35.8%) with 8.8% being non-amplified samples. The F0 mosquitoes were fully susceptible to clothianidin (Supplementary Figure 1). The BusiaUg colony was used for wall bioassay experiments when larval breeding sites were dry in August 2022 and January 2023. The BusiaUg colony was established in 2019 (Oruni et al., 2024) and maintained in the Nagongera, Tororo insectary without insecticide selection. Insecticide resistance characterization showed that the unselected BusiaUg strain was resistant to deltamethrin (Supplementary Figure 1). The Kisumu strain is from western Kenya and is generally susceptible to all insecticides but some tolerance to DDT has been reported (Williams et al., 2019). However, the “Kisumu” strain maintained at the Nagongera insectary was found to be resistant to deltamethrin, permethrin and DDT probably because of colony contamination from wild mosquitoes (Supplementary Figure 1). In 2023, we used a combination of field mosquitoes raised from collected larvae (F0), adult female mosquitoes (F1s) raised from indoor resting collections using an electric Prokopack aspirator from the neighbouring district of Mayuge, and the Kisumu lab strain as in 2022. Similarly, we prioritised the use of females derived from field-collected larvae from the same breeding sites as in 2022 but used adult female (F1) mosquitoes when larval breeding sites were limited.

#### An. funestus mosquitoes

In 2022, *An. funestus* mosquitoes were obtained from indoor resting collections from Busia district and used to raise F1s for experiments. In 2023, most of the collected *An. funestus* mosquitoes from Busia showed very low oviposition rates with very high larval mortality and after several attempts, we opted to collect mosquitoes from Mayuge district (see Figure 1A). Major malaria vectors from Busia and Mayuge districts have been shown to have a similar insecticide resistance profile (Tchouakui et al., 2021).

For raising the F1 progeny, blood-fed indoor-resting mosquitoes were collected and kept for 3-4 days on 10% sugar solution until gravid, and then made to lay eggs by forced egg laying (Morgan et al., 2010). Harvested eggs were then floated in water to hatch into larvae.

Mosquitoes were reared under standard insectary conditions at a temperature of between 24-28°C with 70-90% relative humidity and under an approximate 12:12 photoperiod. Mosquito larvae were reared in larval trays and fed on larval food ad libitum. For *An. funestus*, larval water was changed every three to four days until pupation. Emerged adults were kept in Bugdorm cages while being given 10% sugar solution for 4 -10 days before bioassays.

### WHO wall cone bioassays

Cone bioassays on wall surfaces were conducted following IRS applications in March 2022 and March 2023 (Table 1) using a standardized WHO protocol (WHO, 2006). For the 2022 period, *An. gambiae* exposures were done in July, August, September, and January 2023. However, due to difficulties in rearing, *An. funestus* was only exposed in September 2022. For the 2023 period, *An. gambiae* exposures were done every two months from April 2023 to February 2024. Similarly, due to challenges in rearing sufficient numbers of *An. funestus*, we could only expose them at three time points; September 2023, November 2023 and January 2024. Briefly following the WHO methodology; cones were placed on house walls at three heights, approximately top, middle and bottom. To control for non-insecticide induced mortality a cone was fixed to a wall but with a barrier of paperboard to prevent mosquitoes directly contacting the wall. Three- to ten-day-old female mosquitoes were used for the tests. Field mosquitoes were exposed at the top, middle and bottom while the Kisumu strain and control cone were exposed only at the middle position due to numbers needed (illustrated in Supplementary Figure 2). Mosquitoes were aspirated into the plastic cones in batches of 10 and left exposed on the sprayed surface for 30 minutes. At the end of the 30-minute exposure period, the mosquitoes were carefully collected and transferred to paper cups and provided with 10% sugar solution. One hour after removal from the wall knockdown rates were recorded, and the final mortality was recorded after a 24-hour post-exposure holding period for Actellic and 48 hours for Sumishield, given the delayed mortality expected for slower-acting neonicotinoids (Tchouakui et al., 2022). As a quality control, we performed similar experiments in 15 houses from Busia district that have never received IRS.

### High Performance Liquid Chromatography (HPLC) analysis

#### Sampling insecticide quantity on walls

We sampled insecticide on walls at the same time wall cone bioassays were conducted. In 2022 we were unable to sample at the same time as the cIRS was deployed and sampled from the middle part of the bedroom wall only where four Bostik glue discs were placed around the cone and the insecticide sampled as described in the supplementary section (Supplementary file 1). For each household, one sample was taken at 4-, 5- and 7-months post spraying giving a total of 180 samples comprising of Sumishield sprayed houses (n=51) and unsprayed houses (n=9). In 2023; sampling was done both at the time of spraying and post-spray from both the living room and bedroom. During the spray operations, to enable calculation of extraction efficiency three Whatman filter papers were stuck on the upper, middle and lower part of the house walls in the living room and bedroom. After the spray operation, filter papers were removed, and residual insecticide was sampled from around the same area using four Bostik extra-adhesive glue dots. Extra-adhesive glue dots were used in 2023 to better sample the micro-encapsulated pirimiphos-methyl insecticide during Actellic spraying as previously done (Fuseini et al., 2020). The four glue dots were then carefully removed and stored following the protocol described in the supplementary section (Supplementary file 2). For each household, samples were taken at 0- and 3-month post spraying for Actellic (n=49), Sumishield (n=9) and Unsprayed houses (n=2). The filter papers and glue dots were stored at 4°C before shipping to LSTM for HPLC analysis.

#### HPLC analysis

The analytical standard for pirimiphos-methyl (94.2%) was purchased from Chem Service (USA). The standard for clothianidin (98%) and the internal standard Dicyclohexyl Phthalate (99%) were obtained from Sigma-Aldrich (UK). HPLC grade Acetone (≥99.8%), Acetonitrile (≥99.9%) and deionised water were obtained from Fisher Scientific. Filter paper and glue dot samples were prepared for analysis using a 0.635 cm (½ inch) diameter hole punch. For the filter paper samples, 12 circles were cut out, providing a consistent total surface area of 15.201 cm^2^ for each sample. Glue dot samples consisted of a plastic strip with four glue dots attached to filter paper. Each glue dot was cut out using the hole punch resulting in a total sample surface area of 4.6 cm^2^. Where the glue dot was misshapen and the hole punch could not make a clear incision, scissors were used to cut around the glue dot. Once cut out, the samples were then placed into 10 mL glass extraction tubes. The extraction solution was prepared by dissolving 100 µg/mL of dicyclohexyl phthalate (DCP) in acetone. A 5 mL aliquot of this solution was pipetted into each extraction tube. The tubes were then sealed with tin foil and capped. Filter paper and glue dot samples were sonicated at ambient temperature for 30 minutes and 60 minutes respectively, using an Ultrawave U Series Sonicator. Laboratory optimized protocol requires Glue dot samples to undergo a 60-minute sonication because the residual insecticide is encased within a filter paper and a plastic strip. This causes glue dots to have higher substrate retention than filter papers, which allow insecticide residue to be fully exposed to the extraction solution. Following this, 1 mL of the sonicated solution was transferred into a new 10 mL glass tube, and the solvent was evaporated to dryness using a Techne dri-block (Camlab, UK) under compressed air at 60 °C. Each evaporated sample was resuspended in 1 mL of acetonitrile, and vortexed for 1 minute at 2,500-3,000 rpm. The vortexed samples were poured into 1.5 mL Eppendorf tubes and centrifuged at 13,000 rpm for 15 minutes at ambient temperature. 80 µL of the supernatant was pipetted into a Chromacol 300 µL glass vial (Thermo Scientific, UK), ready for high performance liquid chromatography (HPLC) analysis. HPLC analysis was performed using a Dionex UltiMate 3000 comprising an autosampler, quaternary pump, and variable wavelength detector (Thermo Scientific, UK). A 250 mm × 4.6 mm HPLC column (Thermo Scientific Hypersil Gold C18) was used for both active ingredients (Supplementary Table 1).

Following HPLC analysis, the chromatograms were analysed using Chromeleon 7.3 software. The active ingredient (AI) concentrations were determined using standard curves created from known concentrations of analytical standards for both pirimiphos-methyl and clothianidin. The quantified insecticide levels were corrected against the internal standard (DCP) peak areas, to account for any minor concentration changes in samples during the extraction process. The values were then adjusted for the 1 in 5 dilution.

The concentration of AI in each sample was calculated as follows: (*Where Cs is the concentration of active ingredient in µg per sample, A is the Peak Area on the HPLC, S is the slope value from the standard curve, F is the dilution factor, and P is the DCP Correction factor*)

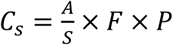

This value was then used to calculate the AI in g/m^2^ (*Where C is the concentration of active ingredient in g/m^2^, Cs is the concentration of active ingredient in µg per sample, and R is the surface area of the sample in cm^2^*)

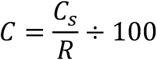

Using a glue dot to extract AI from a wall is not 100% efficient. Therefore, the extraction efficiency of this method must be calculated and factored into final calculations. Extraction efficiency is defined as the proportion of the total residual insecticide on the surface that is recovered by the glue dot. This was calculated by comparing glue dot samples with filter paper samples that were sprayed with the same levels of AI. Since filter papers are expected to retain 100% of the insecticide sprayed onto them, the extraction efficiency of the glue dots can be calculated by comparing their results to those of the filter papers.

Extraction efficiency was calculated as outlined below: (*Where E is extraction efficiency (%), G is the concentration of active ingredient from a glue dot in g/m^2^, and Fp is the concentration of active ingredient from a filter paper in g/m^2^*)

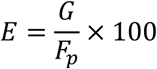

The concentration of AI in the glue dot samples was then corrected for their extraction efficiencies, to estimate the total residual insecticide on the wall, as outlined below: (*Where Gc is the corrected concentration of active ingredient from a glue dot in g/m^2^, E is extraction efficiency (%), and G is the concentration of active ingredient from a glue dot in g/m^2^*)

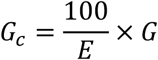

Where filter papers were provided for specific houses, the specific extraction efficiency in that household was used. For households with no filter paper sample provided, the average extraction efficiency was used.

### Temporal profiling of insecticide resistance

In 2023, we assessed insecticide resistance patterns to pirimiphos-methyl and clothianidin using WHO tube assays and bottle assays respectively as per protocol (WHO, 2022b). We used collections from May 2023, August 2023 and November 2023. From the F1 progeny of both *An. gambiae* and *An. funestus,* 3- to 5-day-old female mosquitoes were exposed at different times (5, 15, 30 and 60 minutes) to produce a time response curve for both insecticides and different doses (1µg, 2µg and 4µg per CDC bottle) of clothianidin to assess the dose-response curve. After exposure, 1-hour knockdown was recorded and post-exposure mortality recorded for 5 days for clothianidin to assess delayed mortality and the standard 1 day for pirimiphos-methyl.

### Data analysis

Data were transferred from the data recording sheets into Excel. Knockdown and mortality rates and insecticide residual concentration were analysed and visualised in R/R-studio software (Version 1.4.1106). Test for statistical significance was performed depending on the variables being compared. A comparison of means between knockdown and mortality rates within the same year was performed using the one-way ANOVA test. A comparison of mean knockdown and mortality rates between the years and mosquito species in bioassays was done using a t-test for two independent population means. The wall surfaces were categorized as sub-optimal (concentration less than 0.5 g/m^2^ for Actellic and 0.3g/m^2^ for Sumishield) and compared between months of spray. Similarly, the residue insecticide concentration was averaged by household to determine how many households received adequate or excessive amounts of insecticides.

We used models to establish the relations between mortality and insecticide concentration using R-software (Version 1.4.1106). Pearson’s product-moment correlation was performed to assess the relationship in the residual insecticide concentrations between the different months post-spray. A binomial logistic regression analysis with logit link function was used to examine the relationship between survivorship and residual insecticide concentration as the predictor variable and household as a random effect to address potential clustering. Additionally, the binomial logistic regression also examined the relationship between the occurrence of mortality against various predictors, including insecticide levels, the type of wall material, mosquito strain and months post spray. We then further used the Gaussian regression model with identity logLink to examine the relationship between insecticide levels and different types of wall materials. Lastly, we also ran the linear regression model to examine the relationship between knockdown or mortality rates and residual concentration of insecticides on the walls.

## Results

### Concentration of IRS chemicals sprayed on walls of houses in 2022 and 2023

To test the hypothesis that inadequate insecticide coverage contributed to the resurgence of malaria, we measured the concentration of IRS chemicals sprayed on the walls of houses to determine whether recommended application rates had been achieved. Given we were only able to control for extraction efficiency during the 2023 round, we will discuss these data first. During the 2023 IRS campaign, we took both filter paper and glue dot samples during spraying (0 months) and 3 months post spray from both Sumishield and Actellic sprayed houses.

In Sumishield houses, results from the filter paper extractions showed that none of the houses achieved the target dose (0.3 g/m^2^) (Supplementary Figure 3A) with an average extraction efficiency from the glue dots of 61.27% and limited variation between or within households (variance = 0.01). In Actellic houses, filter paper results showed that over 85% of the households exceeded the optimal levels of insecticide (0.5g/m^2^) (Supplementary Figure 3B) with a low average extraction efficiency of 10.65% and an observed variation between the households (variance = 1.27) but not within the households from different rooms.

In Sumishield houses (n=9) during the 2023 Actellic round, after adjusting for extraction efficiency ∼70% of the surfaces had sub-optimal residual concentration of clothianidin at 0-months post spray. However, when samples were taken at 3-months, only ∼10% of the surfaces had sub-optimal concentrations. More insecticide was detected at 3-months compared to 0-months post spray (*P* < 0.001) (Figure 3A). The cause of this difference is unclear but it could have been non-uniformity in the spray operations since different surfaces were sampled at the different months or technical error during sampling in the field. Furthermore, the sample size of households was too small and these were all mud and pole houses. In Actellic sprayed houses (n=49), only ∼25% of the surfaces had sub-optimal concentrations of pirimiphos-methyl at both 0-months and 3-months post spray. Similarly, more insecticide was detected at 3-months than 0-months (*P* < 0.001) (Figure 3A). We compared the average concentration of insecticides sprayed by wall type, room location and part of the wall. Consistently, “burnt bricks with mud” walls had a significantly higher concentrations of insecticides compared to “mud and poles” walls [(*P* = 0.0078, at 0-months; *P* = 0.0072, at 3-months)] and “burnt bricks with plaster-cement” walls at 3-months (*P* < 0.001) (Figure 3B). There was no significant difference in insecticide concentration between “burnt bricks with plaster-cement” walls and “mud and poles” walls at 0-months (*P* = 0.686) although the “mud and poles” walls had higher concentrations of insecticides at 3-months (*P* = 0.0178) likely due to variations in spray operations. We did not observe any difference in concentrations of insecticides in the living room and bedroom (Figure 3C) and part of the wall (Supplementary Figure 4C) for both Sumishield and Actellic houses.

**Figure. 2:**
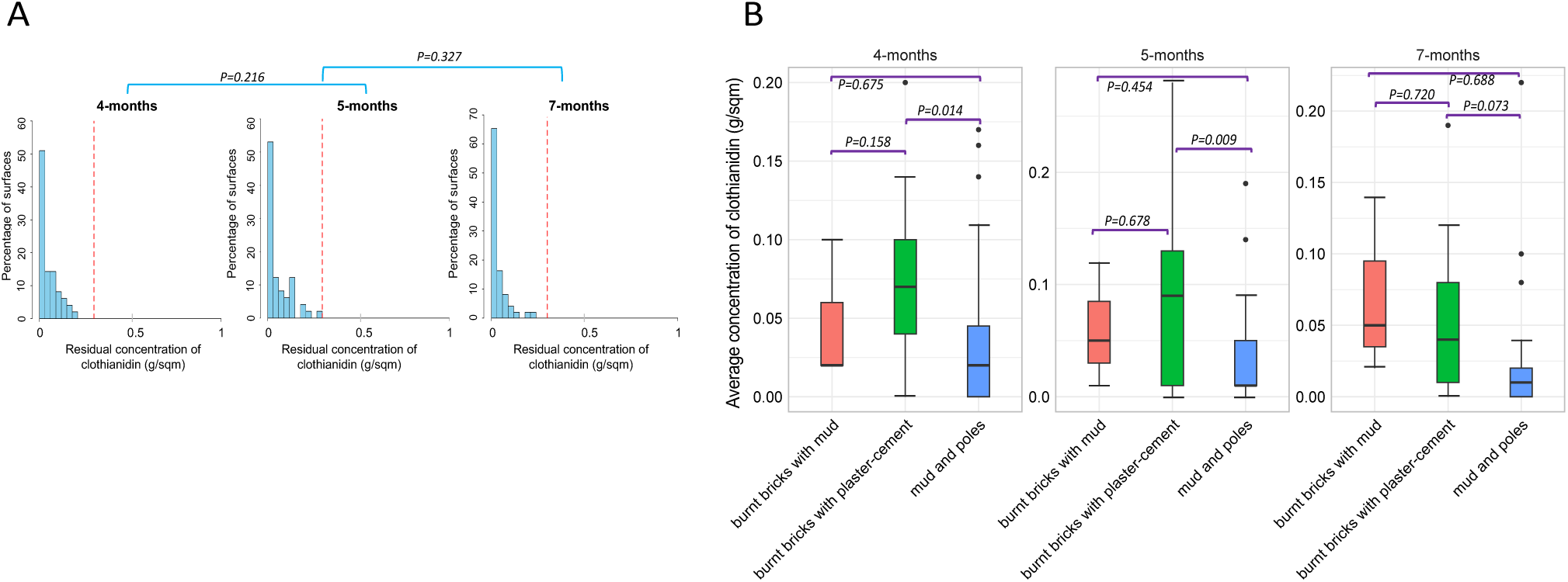
Distribution of IRS insecticide concentrations from household surfaces in 2022 sampled from the middle part of the wall only. A) Histogram of residual concentration of pesticide sampled from 51 households. B) Box plot of the mean concentration of insecticides from different types of walls. The red line represents the reference lines for samples below 0.3 g/m2 (sup-optimal dose).

**Figure. 3:**
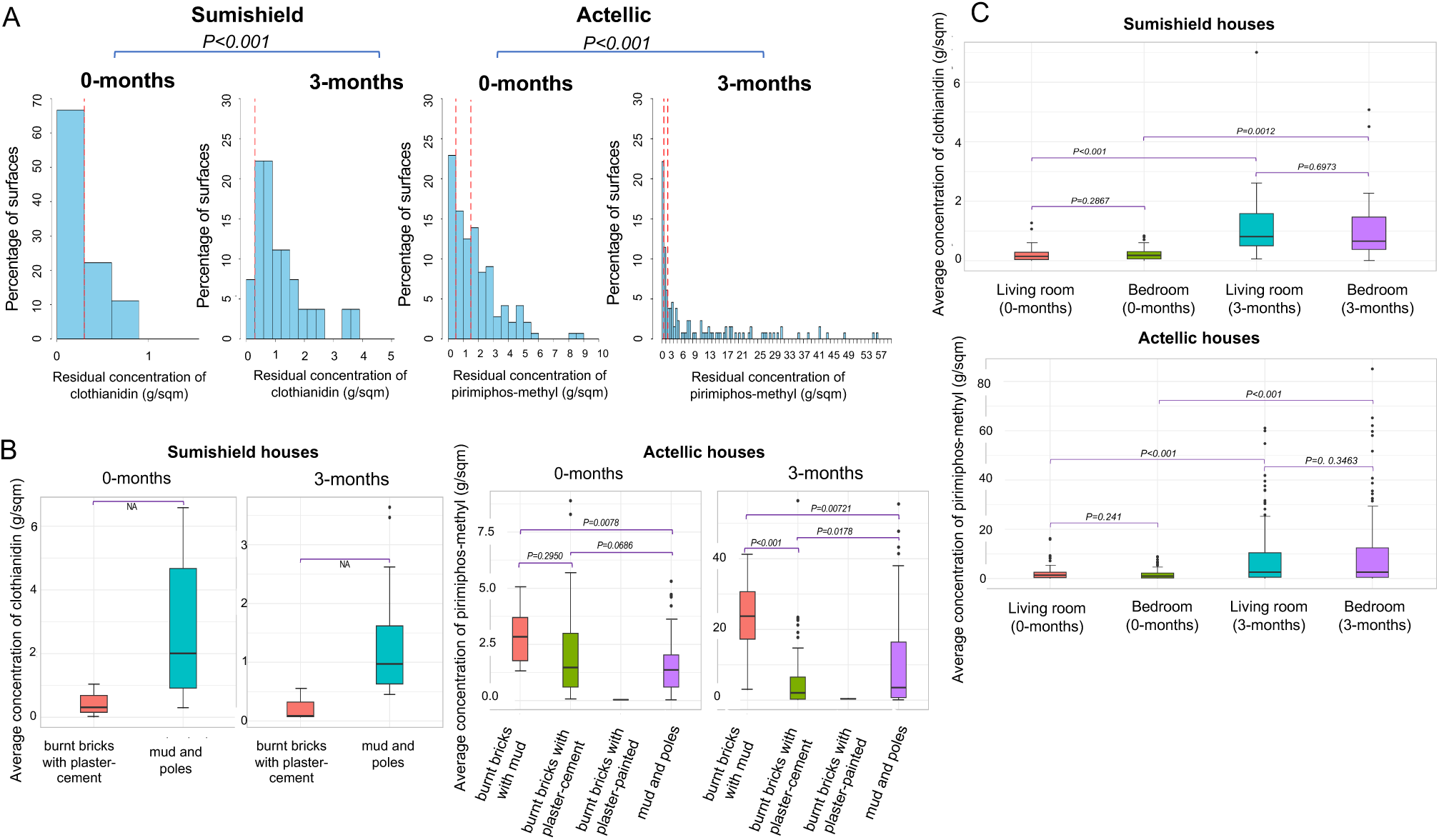
Distribution of IRS insecticide concentrations from household surfaces in 2023. A) Histogram of residual concentration of pesticide sampled from 58 households. Box plot of the average concentration of insecticides from wall types (B) and rooms (C). The dotted red vertical line represents reference lines for samples below 0.3 g/m^2^ (sup-optimal dose for Sumishield) and 0.5 g/m^2^ (sup-optimal dose for Actellic) and 1.5 g/m^2^ where there’s no additional benefit.

During the 2022 Sumishield round, we took samples at 4-, 5-, and 7-months post-spray. One major limitation was that we couldn’t take the samples at the time of spraying and hence couldn’t adjust for extraction efficiency during the HPLC analysis. Our results showed that most of the surfaces had clothianidin, but 50 - 70% of these surfaces had concentrations ≤ 0.05 mg/m^2^. However, at all points of sampling, none of the surfaces achieved the desired 0.3 g/m^2^ optimal level of residual insecticide as per the analysis without adjusting for extraction efficiency. There was no significant difference in clothianidin concentration between the months of sampling post-spray; 4 and 5 months (*P* = 0.216) and 5 and 6 months (*P* = 0.327) (Figure 2A). Additionally, we compared the average concentration of insecticides between wall types at the different months. There was a significant difference in concentrations at 4 months and 5 months but not 7 months. The “burnt bricks with plaster-cement” walls significantly had a higher concentration of insecticide than “mud and poles” walls (4-months; *P* = 0.014, 5-months; *P* = 0.009) but not when compared to “burnt bricks with mud” walls. Similarly, there was no significant difference in insecticide concentration between “mud and poles” walls and “burnt bricks with mud” walls (*P* > 0.05) (Figure 2B).

### Trends in mosquito exposure to IRS chemicals on walls of houses in 2022 and 2023

We assessed the bioefficacy of the insecticides sprayed on the walls against populations of both *An. gambiae* and *An. funestus* in 2022 and 2023. In both years, the control huts in Busia produced no knockdown and mortality (0% ± 0.5) meaning that mortality observed in the wall cone assays in Tororo was unlikely to be due to low quality of reared individuals. In the first year (2022) when Sumishield was used in Tororo district as an IRS formulation, very high mortality rates (80% to 98%) were observed up to 10 months post spray in *An. gambiae* populations including those collected from the wild population (Figure 4A). However, mortality in wild *An. funestus* populations was much lower. When compared in near contemporary assays (6-7 months), knockdown was significantly lower with *An. funestus* compared to *An. gambiae* (7.3% ± 0.4 vs. 50.9% ± 2.5, *P* = 0.00013) as well as 48-hour mortality rates (76.1% ± 3.8 vs. 98.2% ± 4.9, *P* = 0.00043) (Figure 4B).

**Figure. 4:**
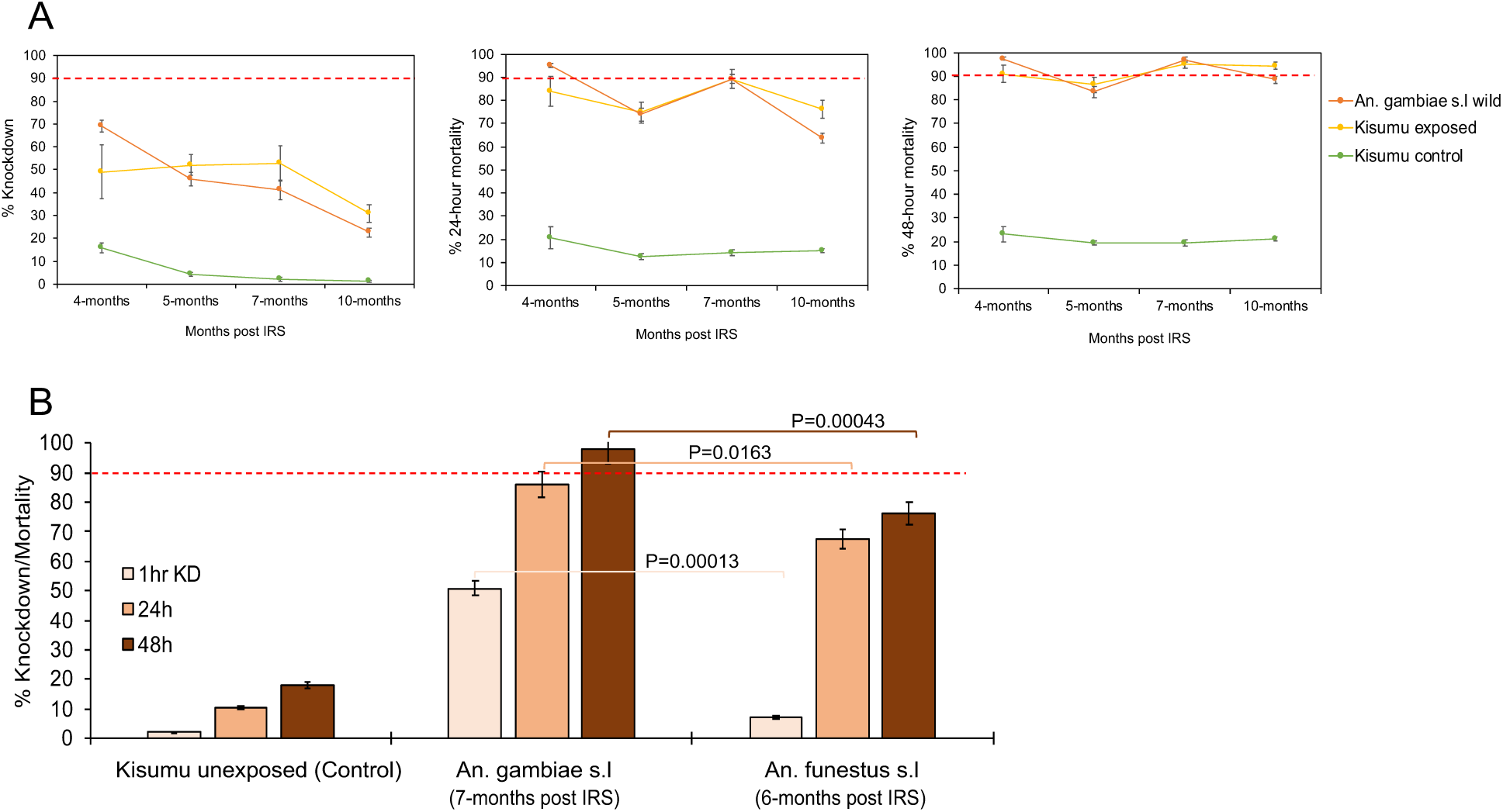
Wall cone assay results for mosquitoes exposed to house walls sprayed with Sumishield at different months post IRS in 2022. A) are the results for *An. gambiae* populations exposed to walls and B) is the result comparing *An. funestus* and *An. gambiae* populations at a single point from the same houses (n=9) where exposure was done. Error bars indicate SEM. The red dotted horizontal line is 90% mortality cut-off, below which is confirmed resistance.

In 2023, two IRS chemicals were sprayed; Sumishield and Actellic. In Actellic sprayed houses, wild *An. gambiae* mosquitoes showed unexpectedly low knockdown (below 40%) and mortality rates (below 80%) from the 1^st^ month up to the 12^th^-month post spray with a general trend of decrease in mortality in the later months (Figure 5A). Compared to the control strain Kisumu, knockdown and mortality rates in wild *An. gambiae* mosquitoes was significantly higher (ANOVA; *F* = 6.85, *df* = 9, *P* = 0.031) meaning that the population of *An. gambiae* might be mildly tolerant to Actellic. When tested against Sumishield sprayed houses, wild population of *An. gambiae* showed higher knockdown (60.2% ± 15.54) and mortality rates (100% ± 0.5) at 1-2 months post spray. Mortality remained above 80% for up to 12 months except at 5-6-months’ time point (59.1% ± 5.75) (Figure 5B). These results indicate that the *An. gambiae* population is largely susceptible to Sumishield.

**Figure. 5:**
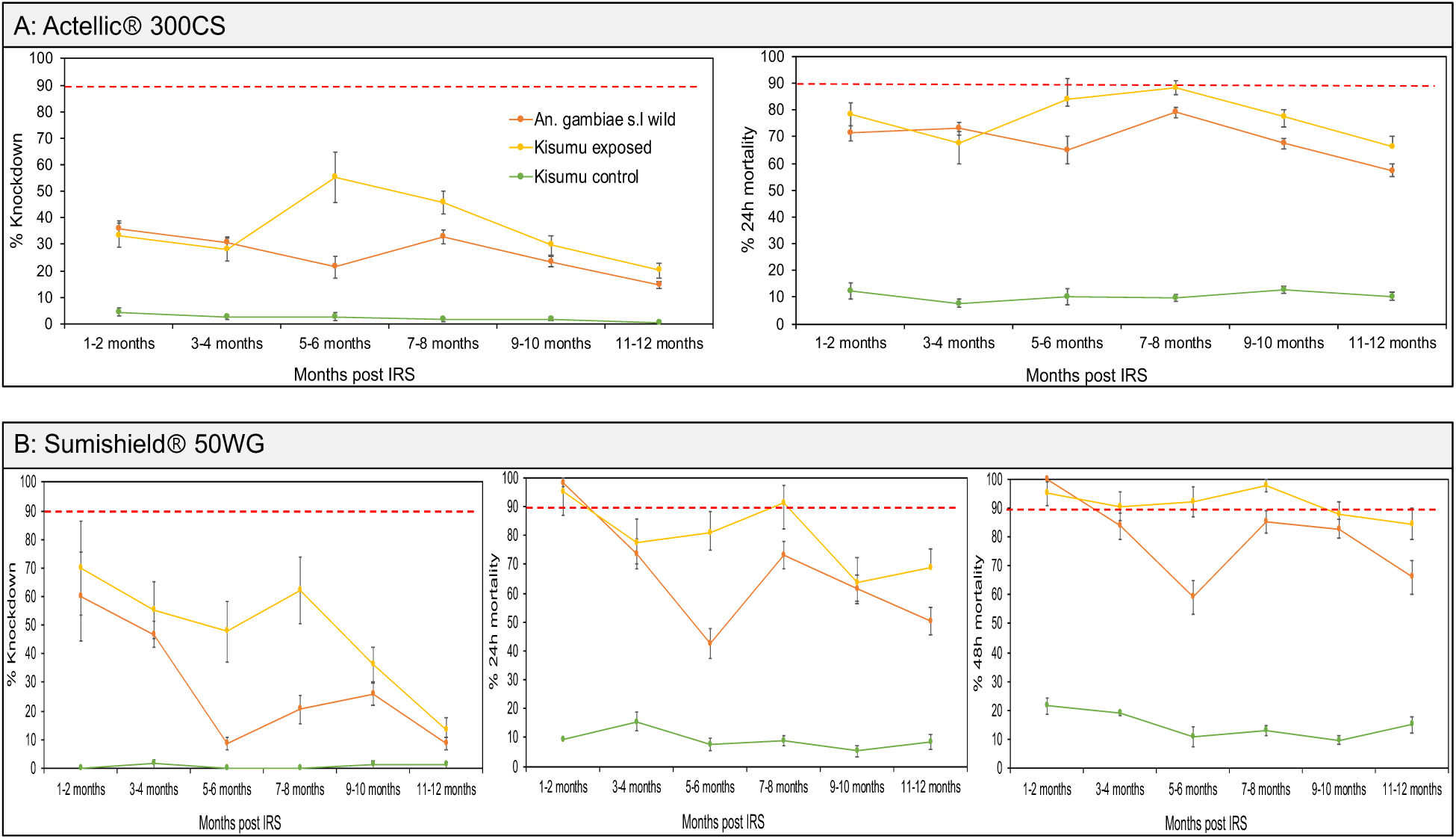
Wall cone assay results for *An. gambiae* mosquitoes exposed to house walls sprayed with Actellic and Sumishield at different months post IRS in 2023. Figure, A) shows the results for walls sprayed with Actellic and B) walls sprayed with Sumishield. Error bars indicate the SEM of the mean. The red dotted horizontal line is 90% mortality cut-off, below which is confirmed resistance.

In contrast, wild *An. funestus* mosquitoes exposed to Actellic, whilst exhibiting the lower knockdown rates (below 40%) had a very high mortality rates up to 6-7-month post spray (92.37% ± 3.56), although a marked decline was observed after this point (8-9 month; 60.29% ± 4.49 and 10-11 month; 51.62% ± 5.1) (Figure 6A). The high susceptibility of *An. funestus* populations to Actellic for up to 7 months contrasted with the Sumishield experiments where mortality rates throughout the months of observation were below 75% compared to the control population Kisumu which had a significantly higher mortality (ANOVA; *F* = 8.82, *df* = 15, *P* = 0.011) (Figure 6B). Hence, the results show that the wild population of *An. funestus* seems to be highly tolerant to Sumishield based on the delayed mortality metric.

**Figure. 6:**
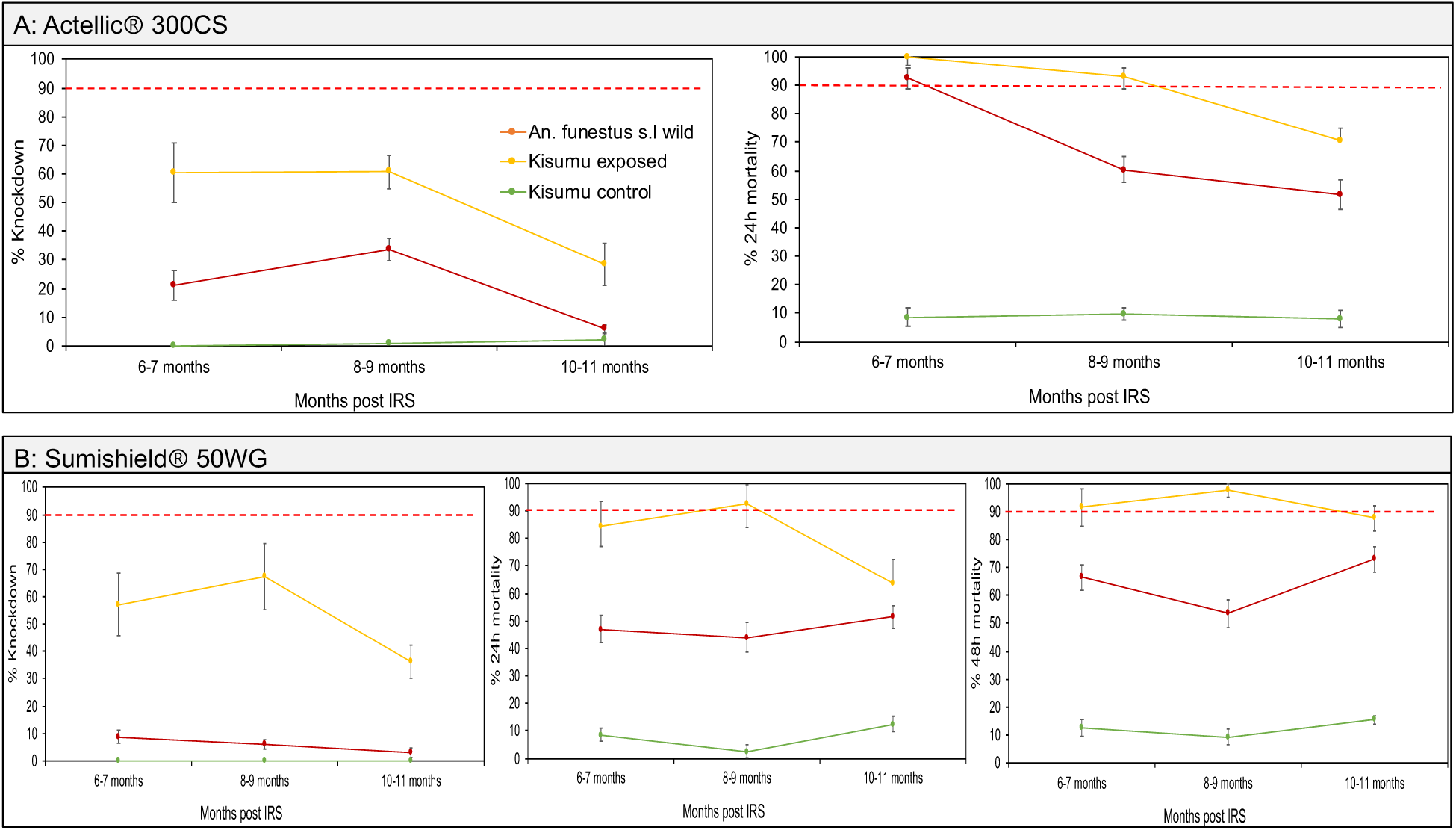
Wall cone assay results for *An. funestus* mosquitoes exposed to house walls sprayed with Actellic and Sumishield at different months post IRS in 2023. Figure, A) shows the results for walls sprayed with Actellic and B) walls sprayed with Sumishield. Error bars indicate the SEM of the mean. The red dotted horizontal line is 90% mortality cut-off, below which is confirmed resistance.

In Sumishield houses, *An. funestus* populations had a significantly lower mortality rates at both 24-hours and 48-hours than *An. gambiae* (Figure 7). When we compared the response to Sumishield in the same vector populations between years; In *An. funestus*, there was a significant difference in mortality rates with higher rates observed in 2022 (average mortality = 72.5%) than in 2023 (average mortality = 64.3%) (*P* = 0.016). In *An. gambiae* populations, there was also a significant difference in mortality rates (48 hours) at all months of comparison with higher rates observed in 2022 than in 2023 (average mortality = 91.5% vs. 79.4%, *P* ≤ 0.003) (Figure 7). The higher mortality rates in 2022 than in 2023 may reflect the greater diversity of wall types studied in 2022 (Table 1) given “mud and pole” walls (8 of 9 walls in 2023) were shown to have high insecticide retention (above).

**Figure. 7:**
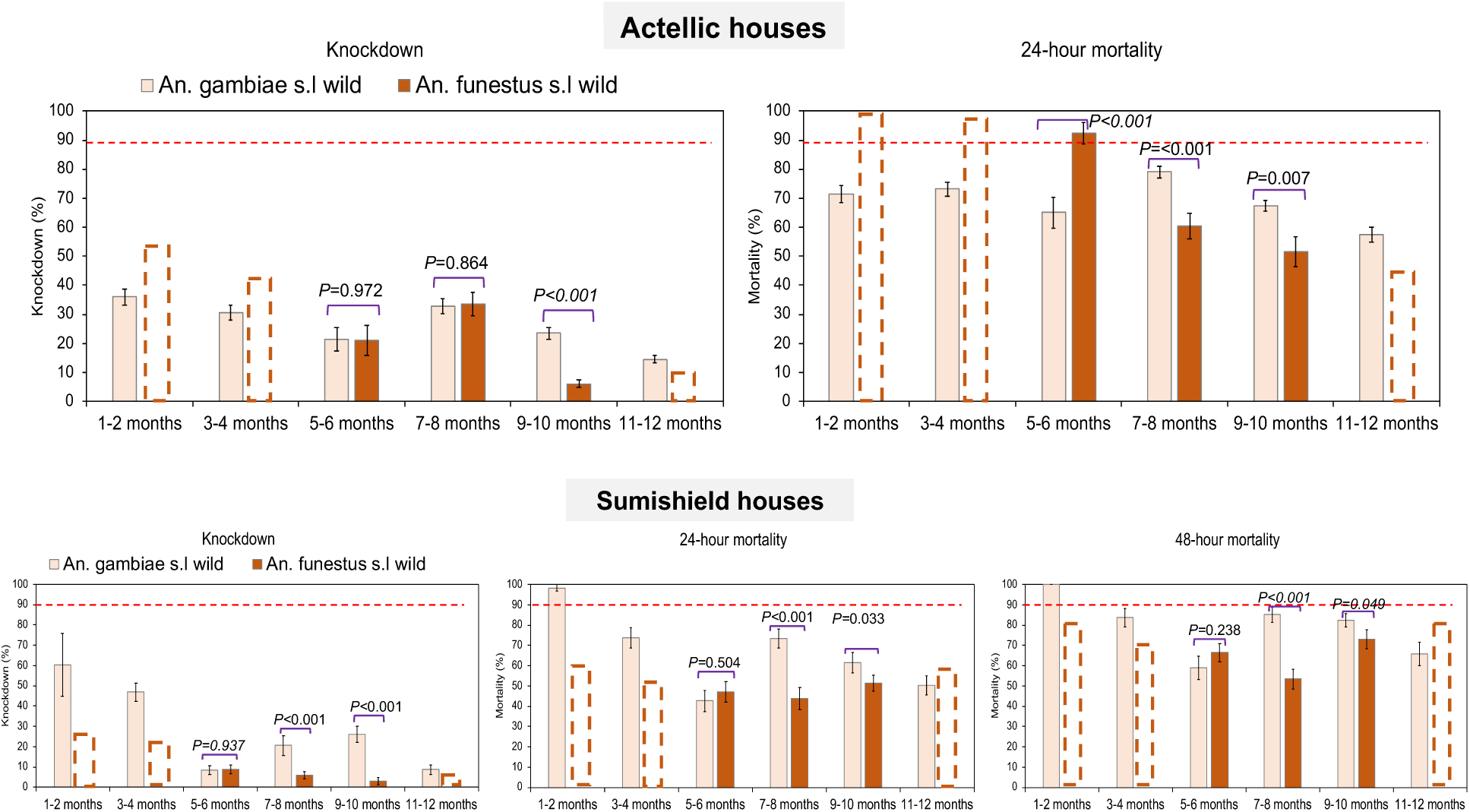
Comparison of wall cone assay mortality between *An. funestus* and *An. gambiae* mosquito populations. The dotted bar-charts indicate the predicted knockdown or mortality based on 3-time points for *An. funestus* for the months that were not done. Error bars represent SEM. The red dotted horizontal line is the 90% mortality cut-off, below which is confirmed resistance.

We compared the differences in spray operation and provide an estimate of variance in mortality by house using data from wild mosquitoes exposed to the different types and parts of walls, selecting only *An. gambiae* populations since it had more data points. In 2022, there was generally a significant difference in knockdown rates at most of the months post-spray by wall type (*P* < 0.05). The “burnt bricks with mud” walls had slightly higher knockdown rates but were very similar to “burnt bricks with plaster-cement” walls while the “mud and poles” walls had significantly lower knockdown rates. However, when we compared mortality rates, there was no significant difference (*P* > 0.05) (Supplementary Figure 3A). Similarly, there was no significant difference in knockdown or mortality rates considering the part of the wall (*P* > 0.05) (Supplementary Figure 3B). It is worth noting that the “burnt bricks with mud” walls had a very small household sample size (n = 3). In 2023, we made the comparison using only Actellic houses which was the main IRS chemical. Considering the type of walls, there was a notable difference in knockdown rates especially in the earlier months (1-6 months post-spray). Again, the “burnt bricks with mud” and “burnt bricks with plaster-cement” walls had significantly higher knockdown rates especially at 1-2 months (*P* < 0.05) and 4-6 months (*P* < 0.001) while “burnt bricks with plaster-painted” and “mud and poles” walls gave lower knockdown rates. Similarly, when we compared mortality rates, “burnt bricks with mud” and “burnt bricks with plaster-cement” walls generally gave higher mortality rates at most of the months post spray compared to “mud and poles” walls (*P* < 0.05), with minor variations including in the “burnt bricks with plaster-painted” walls (Supplementary figure 4A). Contrasting with 2022, there was a significant difference in knockdown and mortality rates by part of the wall in the later months post-spray (7-12 months). The lower part of the wall significantly had lower knockdown rates (*P* < 0.05) except at 11-12 months post spray, and mortality rates (*P <* 0.05) compared to the upper part of the wall that had the highest rates (Supplementary figure 4B).

### Correlation analysis between insecticide wall content and mortality

Table 2 summarizes the correlations between insecticide bioefficacy and key outcome variables, showing varying correlations between residual concentrations at different months, with no to moderate to strong associations in some cases but limited predictive value overall; while binomial logistic regression highlights significant positive associations between insecticide levels and mortality in 2022, Actellic houses in 2023 showed an unexpected negative relationship, and linear regression results suggest positive effects on knockdown and mortality in 2022 but mixed results in 2023, particularly for Actellic houses.

**Table 2:** Summary of correlation correlations analysis between insecticide wall content and mortality using different models.

| Year | IRS House | Model | Variables | Association | Coefficient | SE | p-value |
| --- | --- | --- | --- | --- | --- | --- | --- |
| 2022 | Sumishield | Pearson's product-moment correlation | 4 months post spray versus 5 months post spray | weak positive correlation | $r = 0.07$ | - | 0.0752 |
| 2022 | Sumishield | Pearson's product-moment correlation | 5 months post spray versus 7 months post spray | moderate positive correlation | $r = 0.25$ | - | <0.001 |
| 2022 | Sumishield | Pearson's product-moment correlation | 4 months post spray versus 7 months post spray | moderate positive correlation | $r = 0.35$ | - | <0.001 |
| 2023 | Actellic | Pearson's product-moment correlation | 0 months post spray versus 3 months spray | moderate positive correlation | $r = 0.246$ | - | < 0.001 |
| 2023 | Sumishield | Pearson's product-moment correlation | 0 months post spray versus 3 months spray | strong positive correlation | $r = 0.73$ | - | < 0.001 |
| 2022 | Sumishield | Binomial logistic regression analysis | Residue insecticide levels and the log odds of mortality versus survival | positive association | $\beta = 0.016$ | 0.002 | < 0.001 |
| 2023 | Sumishield | Binomial logistic regression analysis | Residue insecticide levels and the log odds of mortality versus survival | no association | $\beta = 0.090$ | 1.088 | 0.934 |
| 2023 | Actellic | Binomial logistic regression analysis | Residue insecticide levels and the log odds of mortality versus survival | negative association | $\beta = -0.026$ | 0.01 | 0.013 |
| 2022 | Sumishield | Linear regression model | insecticide levels and knockdown rates | positive association | $\beta = 0.214$ | 0.035 | 0.001 |
| 2022 | Sumishield | Linear regression model | insecticide levels and mortality rates | positive association | $\beta = 0.008$ | 0.003 | 0.01 |
| 2023 | Sumishield | Linear regression model | insecticide levels and knockdown rates | no association | $\beta = 1.280$ | 1.029 | 0.223 |
| 2023 | Sumishield | Linear regression model | insecticide levels and mortality rates | no association | $\beta = -4.097$ | 6.029 | 0.502 |
| 2023 | Actellic | Linear regression model | insecticide levels and knockdown rates | negative association | $\beta = -0.087$ | 0.027 | 0.001 |
| 2023 | Actellic | Linear regression model | insecticide levels and mortality rates | no association | $\beta = -0.214$ | 0.211 | 0.313 |

### Temporal monitoring of insecticide resistance to clothianidin and pirimiphos-methyl

We used bioassays to assess the resistance profile of both populations of *An. gambiae* and *An. funestus* to clothianidin and pirimiphos methyl. The results revealed significant variations in the resistance profile temporally.

For clothianidin time exposure assays (Figure 8A); In May 2023, *An. funestus* showed very low knockdown rates, with no knockdown recorded at lower exposure times (5-min, 10-min, 15-min) and rates below 20% recorded in 30-mins (20% ± 0.5) and 60-mins (16.7% ± 0.5). The mortality recorded after 5 days showed lower rates (30 - 50%) for 5-mins, 10-mins, 15-mins and 30-mins and 83.3% for 60-mins, the standard exposure time. However, in the same period, *An. gambiae* had a significantly higher knockdown (35 - 55%) (t = 5.7, *P* = 0.0006) and mortality (78 - 98%) (t = 2.9, *P* = 0.0132) compared to *An. funestus*. In the later months, higher knockdown rates; in August (50 - 75%), November (60 - 85%) and mortality rates in August (95 - 100%), and November (90 - 98%) were recorded in *An. funestus* populations. Likewise, in *An. gambiae* populations higher knockdown rates were recorded in August (85 - 100%) and November (90 - 95%) together with very high mortality rates; in August (80-100%) and November (95 - 100%). The knockdown rates recorded were significantly lower in *An. funestus* compared to *An. gambiae* [August (t = 6.01, *P* = 0.000477) and November (t = 2.80288, *P* = 0.015523)] but there was no significant difference in mortality rates [August (t = 0.68, *P* = 0.2623) and November (t = 1.84, *P* = 0.0577)].

**Figure. 8:**
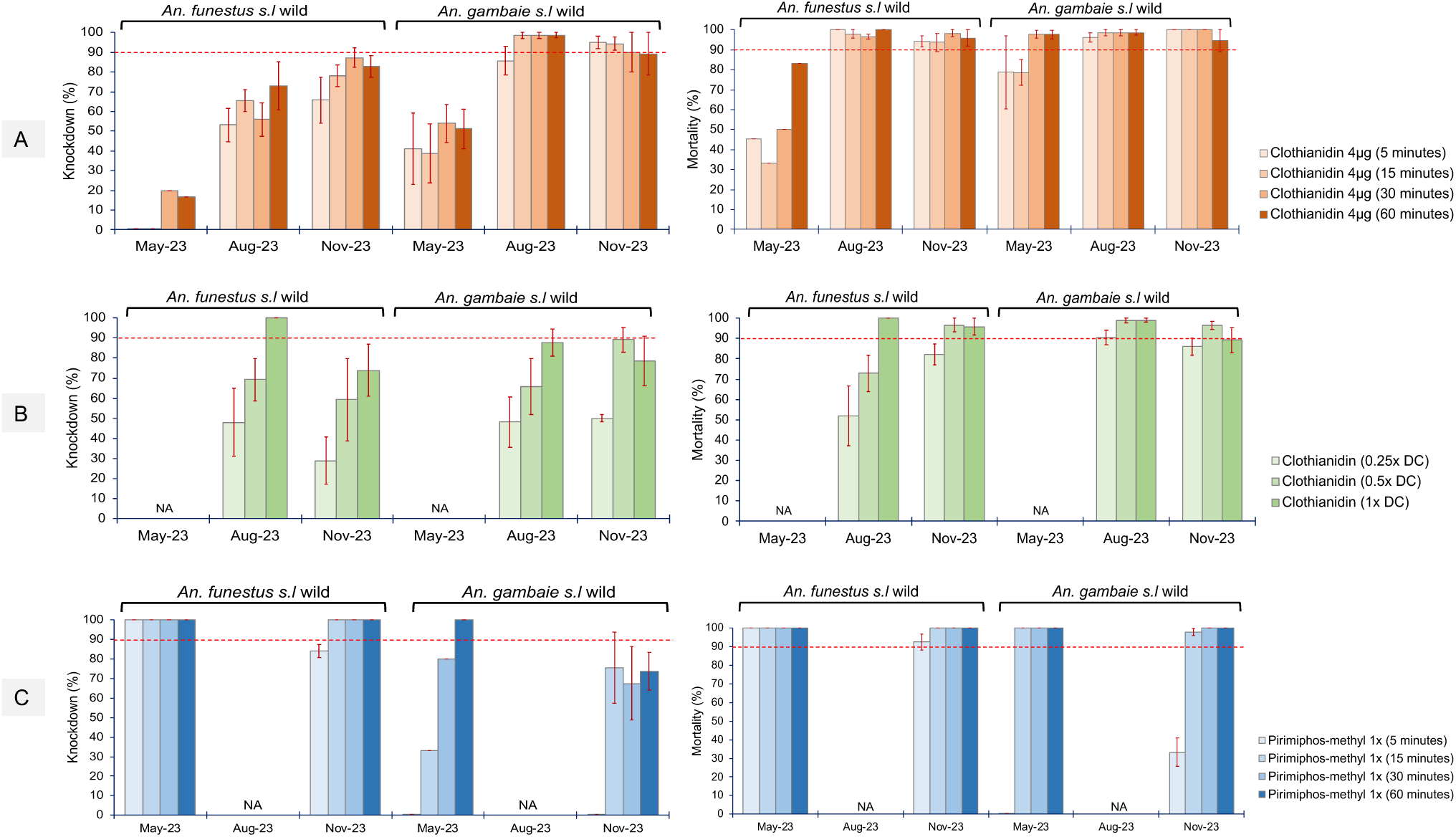
Temporal monitoring of mortality in *An. funestus* and *An. gambiae* mosquito populations using bioassays. A) Time exposure assay for clothianidin, B) Dose-response exposure assay for clothianidin at different diagnostic doses (DC) and C) Time exposure assay for pirimiphos-methyl. Error bars represent SEM. The red dotted horizontal line is 90% mortality cut-off, below which is confirmed resistance.

For clothianidin dose-response assays (Figure 8B); Although we observed slightly lower knockdown rates in *An. funestus* [August (45-100%), November (25 - 75%)], it was not significantly different from the rates recorded in *An. gambiae* populations [August (45 - 90%), November (50 - 90%)] [August (t = 0.27, *P* = 0.4) and November (t = 1.05, *P* = 0.176)] and similar results were observed in the mortality rates.

For pirimiphos-methyl time exposure assays (Figure 8C); *An. funestus* showed significantly higher knockdown rates [May (100%), November (80 - 100%)] compared to *An. gambiae* populations [May (0- 100%), November (0 - 80%)] [May (t = 2.07, *P* = 0.042282), November (t = 2.25, *P* = 0.033)]. Nonetheless, both vector populations were susceptible to pirimiphos-methyl with no significant difference in mortality (t = 1.44, *P* = 0.0853), although *An. gambiae* populations had very low mortality rates (0 – 35%) at 5-min exposure.

## Discussion

Uganda has largely had commendable success using IRS campaigns for malaria control especially in very high transmission districts (Namuganga et al., 2021) as a supplement to the large-scale distribution of LLINs like the PBO net distribution in 2017 (Staedke et al., 2020). A dramatic malaria resurgence occurred in 2020-2022 which coincided with the introduction of entirely new, clothianidin-based formulations Fludora Fusion and Sumishield (Epstein, Maiteki-Sebuguzi, et al., 2022). During the resurgence, malaria incidence increased by more than 8-fold (0.36 vs. 2.97 episodes per person year, *P* < 0.0001) from 2021 to 2022 when Fludora Fusion and Sumishield were used respectively (Kamya et al., 2024). We sought to specifically investigate the vector dynamics that might have at least in part contributed to this resurgence by assessing the bioefficacy of IRS insecticides (Sumishield and Actellic) against wild populations of two major vectors; *An. funestus* and *An. gambiae* during the 2022 and 2023 IRS rounds. Firstly, we measured if the optimal concentration and type of insecticide were sprayed on the walls of houses and secondly, we assessed the bioefficacy of both Sumishield and Actellic against wild resistant populations of *An. gambiae* and *An. funestus*. We detected clothianidin residues in all of the sprayed houses, confirming that the correct chemical was applied. Similarly, we detected pirimiphos-methyl in all the sprayed houses in 2023 during the Actellic round. When we adjusted for extraction efficiency in 2023, most of the sprayed surfaces (∼70%) had the target concentration of clothianidin although there were significant variations between rounds of sampling. Likewise, during the Actellic round, most of the sprayed surfaces (∼75%) had above the target concentration of pirimiphos-methyl. However, we did notice significant variations in insecticide concentration within the houses considering the wall position. Furthermore, taking Actellic as a comparative example, the levels of pirimiphos-methyl insecticide detected during the IRS round were significantly higher than what would be expected if spray operations are done correctly. A similar study conducted in Bioko Island aimed at improving spray operations, using the same protocol as this study, reported relatively lower but normal levels of pirimiphos-methyl on the walls (Fuseini et al., 2020). Whilst we did not observe any significant difference in insecticide concentrations between different rooms, there were significant differences in insecticide levels on the different wall surfaces like in mud and pole walls which had the lowest concentration of insecticides since they are more porous and therefore insecticide retention was expected to be lower. Indeed, the bioefficacy observed in mud and pole walls in this study was lower than observed in burnt bricks and plaster or cement or painted especially in the earlier months post-spraying. Comparatively, unlike Actellic that is known to retain bioefficacy between 6-12 months (Ibrahim et al., 2017; Oxborough et al., 2014) Sumishield showed extended bioefficacy up to almost 24 months post spray in unsprayed houses against *An. gambiae* population. This is in concordance with a recent study that showed that Sumishield residual activity can last up to 18 months on different wall materials against different mosquito vectors (Lees et al., 2022) highlighting the huge potential of clothianidin-based formulations in areas where vectors are susceptible. Prior to the rollout of clothianidin-based formulations for IRS, a cross-sectional study in 2016-17 established that malaria vectors were largely susceptible to clothianidin-based formulations across sub-Saharan Africa (Oxborough et al., 2019). It is possible that clothianidin resistance or tolerance in *An. funestus* could have developed before this survey since it was not previously detected by Oxborough et al in Tororo but low mortality (<80%) was observed in 2020 in Busia and Mayuge districts using 13.2 mg/m^2^ clothianidin (Tchouakui et al., 2021). In Cameroon, *An. gambiae s.s* populations resistant to clothianidin were discovered in vegetable farms in 2019-20 where usage of clothianidin-based products for pest control was reported (Fouet et al., 2024). The Cameroon study probably features the only well-documented report of clothianidin resistance in Africa but highlights the role of pesticide use in agriculture in producing insecticide-resistant vector populations. Conversely, our study found that *An. gambiae* population was largely susceptible to clothianidin similar to what has been reported in 2017 (Oxborough et al., 2019). However, *An. funestus* mosquitoes were tolerant to clothianidin although with temporal variation in susceptibility, perhaps reflecting changing population dynamics and seasonal selective pressures (Tene-Fossog et al., 2022). In studies from the same area in Uganda, *An. funestus* had reduced mortality to clothianidin in February 2020 (Tchouakui et al., 2021) but very high mortality in October 2021 (Assatse et al., 2023). This pattern is consistent with our study when low mortality was observed in May 2023 but higher mortalities in the months of August and November that may suggest seasonal variations in resistance patterns.

Under field conditions, *An. funestus* consistently showed lower mortality to Sumishield sprayed walls in 2022 and 2023 than Actellic sprayed walls. The contrasting responses of *An. gambiae* and *An. funestus* to clothianidin, provides a compelling explanation for the increase in *An. funestus* abundance compared to *An. gambiae* during the malaria resurgence (Kamya et al., 2024). Similarly, after Actellic was sprayed in 2023 there was a decrease in both *An. funestus* abundance and malaria cases, with a slight increase in abundance of *An. arabiensis* (Kamya et al., 2024). Our study also provides evidence that the reversion was a result of differing insecticide susceptibility in the main malaria vectors given *An. funestus* was susceptible to pirimiphos-methyl while *An. gambiae* (where the majority were *An. arabiensis*) was mildly tolerant to pirimiphos-methyl. Our findings suggest that the dramatic malaria resurgence that occurred in Uganda after the switch to clothianidin-based formulations was driven by *An. funestus*. Unlike *An. gambiae*, *An. funestus* mostly breeds throughout the year and is known to sustain malaria transmission during dry seasons (Doucoure et al., 2020; Menze et al., 2016). Moreover, a recent review has shown that *An. funestus* is now the dominant vector in most localities in east and southern Africa which was likely due to the scale-up of LLINs between 2010 and 2020 (Msugupakulya et al., 2023). The exact mechanisms mediating *An. funestus* tolerance to clothianidin are not known but this is an active area of investigation.

## Conclusion

Insecticide resistance monitoring in malaria vectors is a vital activity that should inform the rollout of control interventions. Notwithstanding the variations observed in spray operations, we are confident that we have retrospectively implicated clothianidin resistance in the major malaria vector *An. funestus* as the likely driver of IRS control failure and it would be preferable for susceptibility testing including wall-based bioassays to be conducted in advance of class switches. Clearly, given the evidence of efficacy of clothianidin-based IRS in other locations like Benin and Cameroon (Ngufor et al., 2017; Oxborough et al., 2019; Thiomela et al., 2024), local testing is required. This is largely due to the significant differences and complexities in vector populations between regions or even countries. The rapid response of the NMCP to re-introduce Actellic was crucial situation for the short-term alleviation of the problem but signals of possible development of tolerance/resistance to pirimiphos methyl in *An. gambiae* populations in both Uganda and the wider region (Lucas et al., 2023, 2024) should not be ignored. Additionally, it may also be prudent as a country to have additional training and refresher courses for spray operators to harmonise the spraying operations to avoid variations that could produce inadvertent consequences. However, the results from this study are location specific and should not be extrapolated too widely given clothianidin-based interventions are effective in other regions largely against *An. gambiae s.l* populations (Ngufor et al., 2017; Ngwej et al., 2019; Oxborough et al., 2019). One clear advantage of clothianidin-based formulation is prolonged residual activity as seen in this and other studies (Lees et al., 2022; Ngufor et al., 2017; Ngwej et al., 2019) albeit with significant variations between wall surfaces exist which is common with most IRS insecticides.

## Data availability

All datasets generated or analysed during this study are included in this published article and its supplementary files.

## Supporting information

Protocol for sampling walls for Fludora Fusion/Sumishield

Protocol for sampling walls for Actellic

## Acknowledgements

We thank all the PRISM study team members for their untiring support during this study and the overall commendable effort and support during the PRISM studies, particularly the Tororo and Busia district administrations. We are very grateful to the study participants who participated in this study. We are grateful to the administration of LSTM, IDRC and UVRI for the additional support during sample shipment, fieldwork and experiments.

## Author contributions statement

**Conceptualization:** Ambrose Oruni, Charles S. Wondji, Grant Dorsey, Martin J Donnelly.

**Data curation:** Ambrose Oruni, Adiga James, Geoffrey Otto, Patrick Kygamba, Daniel Ayo, Jackson Asiimwe, Joseph Okoth, Maato Zedi, Kyle J Walker, Ashlee Braithwaite.

**Formal analysis:** Ambrose Oruni, Kyle J Walker, Ashlee Braithwaite, Martin J Donnelly.

**Funding acquisition:** Ambrose Oruni, Paul Krezanoski, Martin J Donnelly, Grant Dorsey.

**Methodology:** Ambrose Oruni, Hanafy M Ismail, Mark J I Paine, Melissa D Conrad, Martin J Donnelly.

**Supervision:** Emmanuel Arinaitwe, John Rek, Hanafy M Ismail, Mark J I Paine, Melissa D Conrad, Jonathan Kayondo, Charles S. Wondji, Teun Bousema, Moses R. Kamya, Paul Krezanoski, Grant Dorsey, Martin J Donnelly.

**Writing – original draft:** Ambrose Oruni, Kyle J Walker, Ashlee Braithwaite, Martin J Donnelly.

**Writing – review & editing:** all authors.

## Funding

This study was funded by the IDRC/ICEMR Special Project grant awarded to Ambrose Oruni funded by a parent grant from the US National Institutes of Health (NIH) as part of the International Centers of Excellence in Malaria Research (ICEMR) program (U19AI089674). Additional support was given by was provided by an NIH Career Mentored Award (K23AI139364) awarded to Paul Krezanoski. The funders played no role in the design of the study; in the collection, analyses, and interpretation of data; in the writing of the manuscript; or in the decision to submit the manuscript for publication.

## Conflict of interest

The authors declare no conflict of interest.

## Notes

### Competing Interest Statement

The authors have declared no competing interest.

