## Supplementary Figures for "Significant variations in tolerance to clothianidin and pirimiphos-methyl in *Anopheles gambiae* and *Anopheles funestus* populations during a dramatic malaria resurgence despite sustained indoor residual spraying in Uganda"

### Supplementary material

**Supplementary table. 1:** *Shows HPLC methods for different active ingredients.*

| | Active Ingredient | |
|---|---|---|
| | **Pirimiphos Methyl** | **Clothianidin** |
| **Injection Volume** | 20 µL | 20 µL |
| **Mobile Phase** | 70:30 Acetonitrile: Water | 93:7 Acetonitrile: Water with 0.1% phosphoric acid |
| **Flow Rate** | 1 mL/min | 1 mL/min |
| **Run Time** | 22 min | 9 min |
| **Wavelength** | 232 nm | 232 nm |

**Supplementary Figure. 1:** Phenotypic resistance characterisation of BusiaUg and Kisumu lab strains to pyrethroids (PY) – deltamethrin and permethrin and an organochlorine (OC) – DDT, and response of wild F0 mosquitoes to clothianidin + deltamethrin (CTD+DM) and clothianidin (CTD) only.

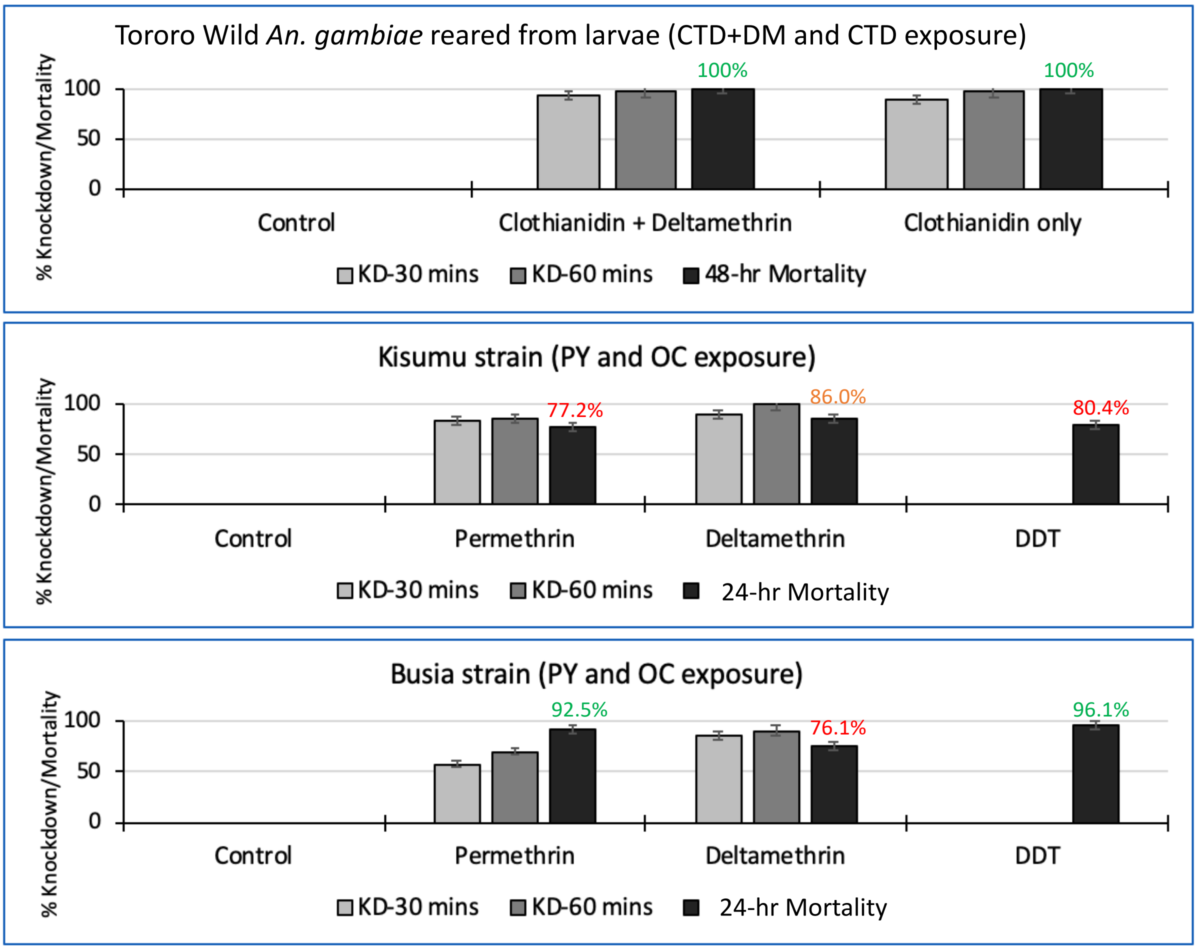

**Supplementary Figure. 2:** A cartoon showing the setup of cones on the walls of houses during exposure. The exposed field mosquitoes are labelled as (Ef), the exposed Kisumu strain (Ek) and the unexposed control cone (C).

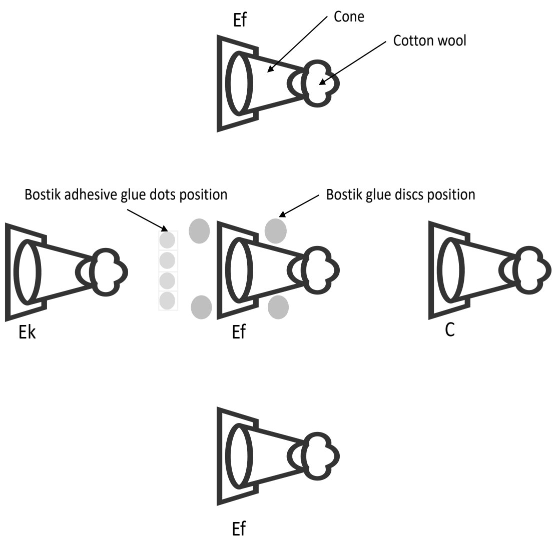


**Supplementary Figure. 3:** Residual concentration of insecticides sampled during spraying (0-months). A) Sumishield (clothianidin) and B) Actellic (pirimiphos-methyl).

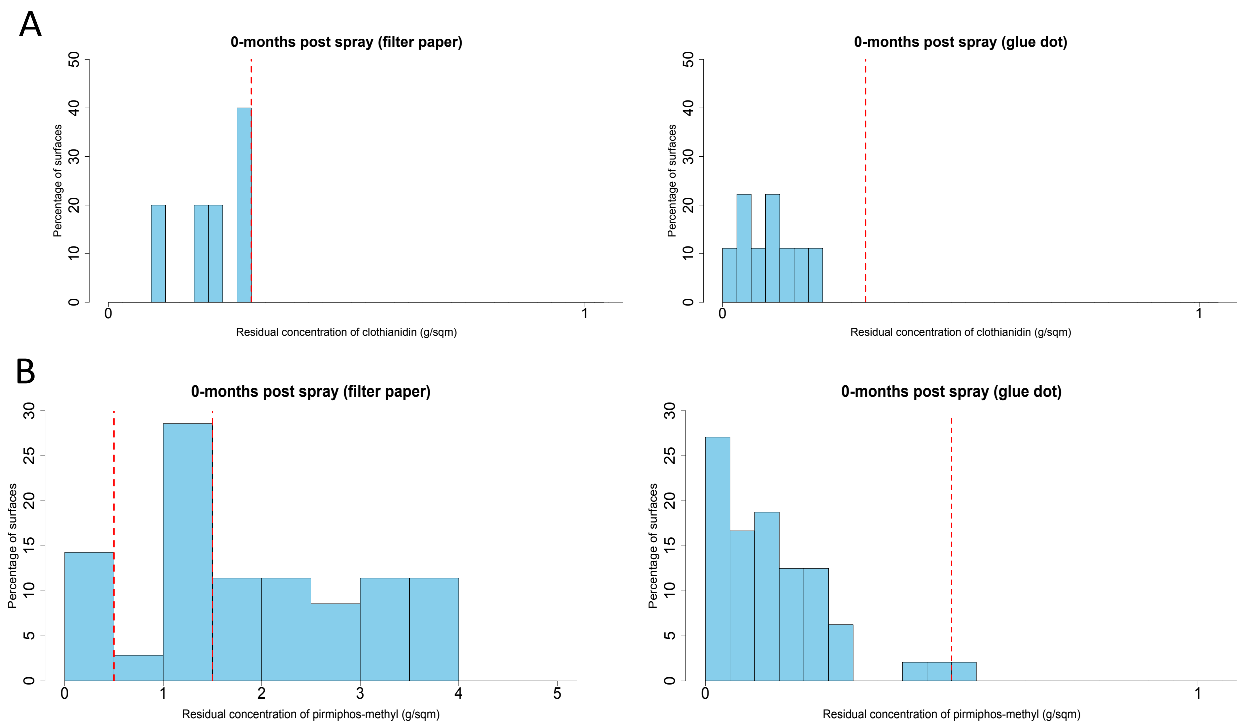


**Supplementary Figure. 4:** Box plot for mortality rates of exposed mosquitoes from wall cone assays in 2022 grouped by wall types (A) and part of the wall (B). Error bars represent SEM. The red dotted horizontal line is

90% mortality cut-off, below which is confirmed resistance. The level of significance is indicated by asterisks (*<0.05, **<0.01, ***<0.001) and NS meaning "not significant".

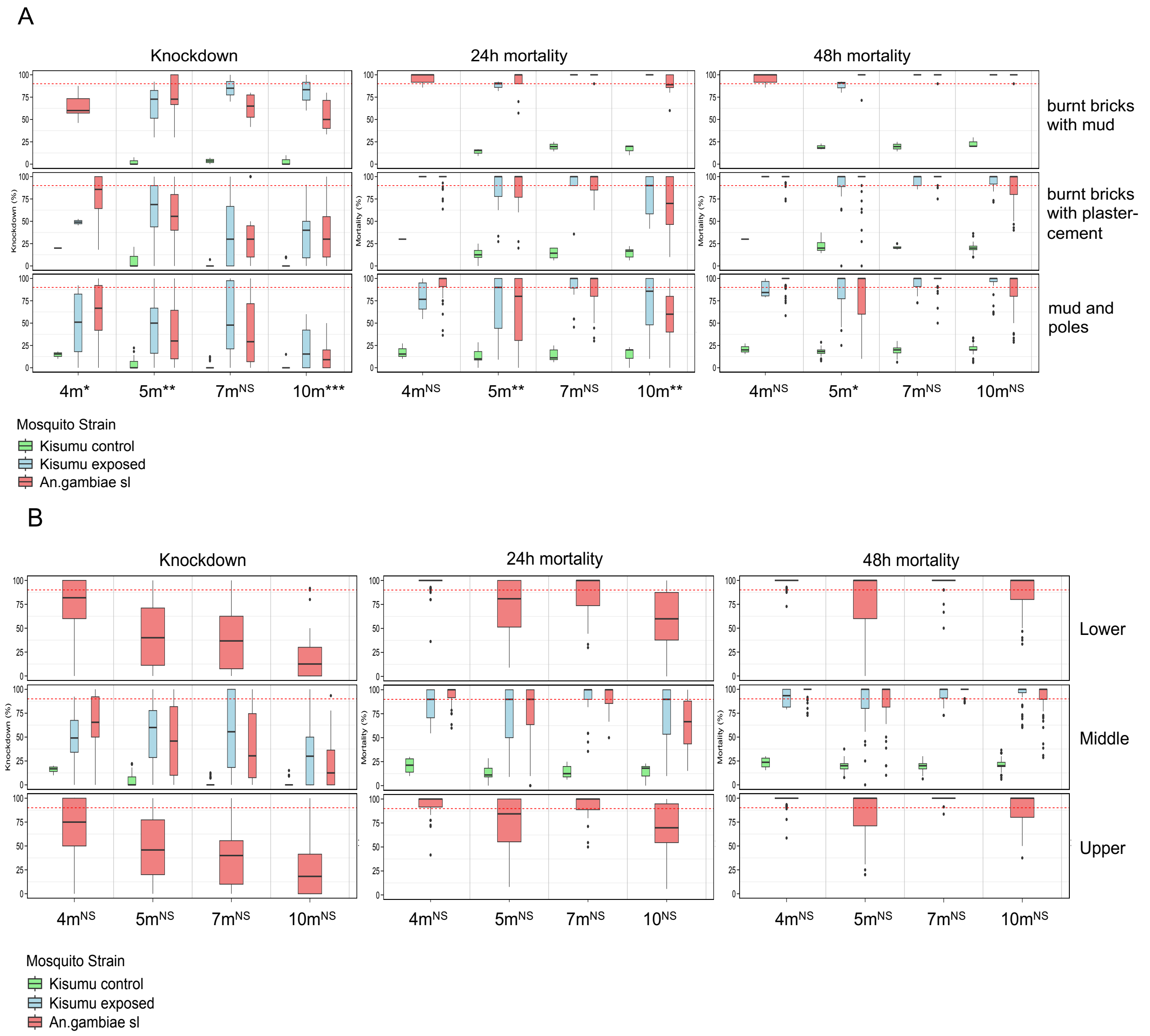


**Supplementary Figure. 5:** Box plot for mortality rates of only *An. gambiae* exposed mosquitoes from wall cone assays in 2023 grouped by wall types (A) and part of the wall (B). C) is the average concentration of IRS insecticides in Sumishield and Actellic houses grouped by part of the wall. Error bars represent SEM. The red dotted horizontal line is 90% mortality cut-off, below which is confirmed resistance. The level of significance is indicated by asterisks (*<0.05, **<0.01, ***<0.001) and NS meaning "not significant".

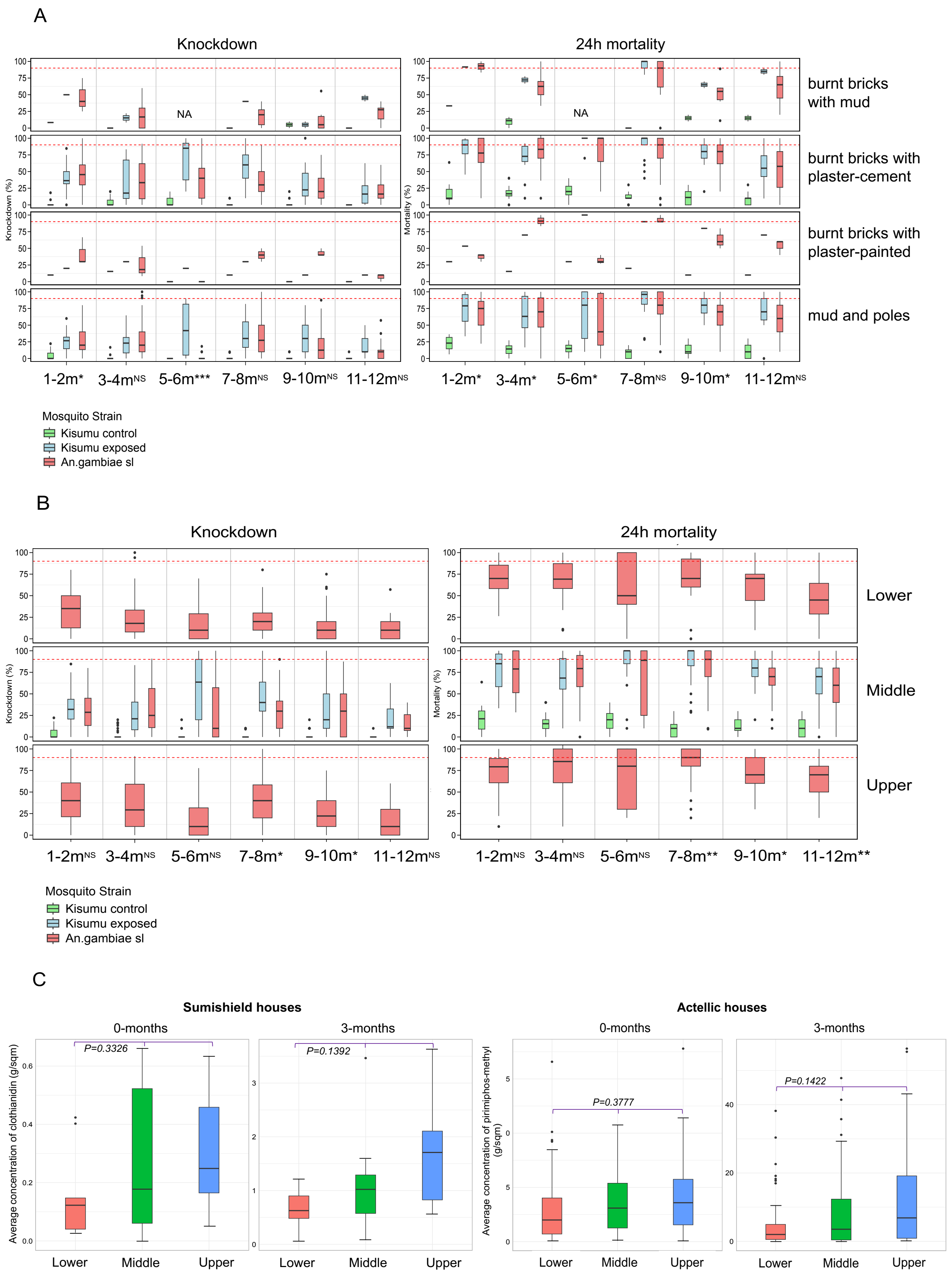
A
Knockdown
24h mortality
Knockdown (%)
Mortality (%)
NA
NA
burnt bricks with mud
burnt bricks with plaster-cement
burnt bricks with plaster-painted
mud and poles
1-2m* 3-4mNS 5-6m*** 7-8mNS 9-10mNS 11-12mNS
1-2m* 3-4m* 5-6m* 7-8mNS 9-10m* 11-12mNS
Mosquito Strain
Kisumu control
Kisumu exposed
An.gambiae sl
B
Knockdown
24h mortality
Knockdown (%)
Mortality (%)
Lower
Middle
Upper
1-2mNS 3-4mNS 5-6mNS 7-8m* 9-10m* 11-12mNS
1-2mNS 3-4mNS 5-6mNS 7-8m** 9-10m* 11-12m**
Mosquito Strain
Kisumu control
Kisumu exposed
An.gambiae sl
C
Sumishield houses
Actellic houses
0-months
3-months
0-months
3-months
Average concentration of clothianidin (g/sqm)
Average concentration of pirimiphos-methyl (g/sqm)
P=0.3326
P=0.1392
P=0.3777
P=0.1422
Lower Middle Upper
Lower Middle Upper
Lower Middle Upper
Lower Middle Upper

**Supplementary Figure. 6:** Bar plot for mortality rates of only wild *An. gambiae* mosquitoes exposed to non-sprayed walls in 2022 and 2023 with a table showing the average concentration of clothianidin detected in those houses.

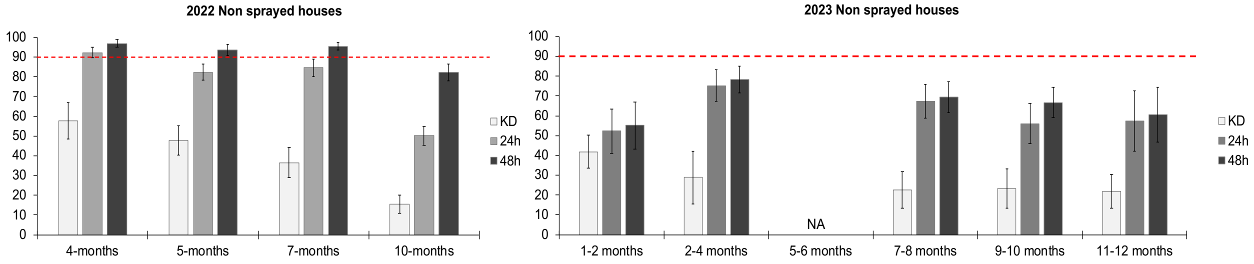


| | 2022 (n=9) | 2023 (n=2) |
|---|---|---|
| Average clothianidin concentration (g/sqm) | 0.0184 | 0.4237 |
| Average pirimiphos-methyl concentration (g/sqm) | NA | NA |
