## Supplementary material for "Significant variations in tolerance to clothianidin and pirimiphos-methyl in *Anopheles gambiae* and *Anopheles funestus* populations during a dramatic malaria resurgence despite sustained indoor residual spraying in Uganda": Protocol for sampling walls for Fludora Fusion/Sumishield

*Sampling protocol*

<u>**Sampling insecticide on walls for Fludora®Fusion/Sumishield® 50WG sprayed surfaces:**</u>

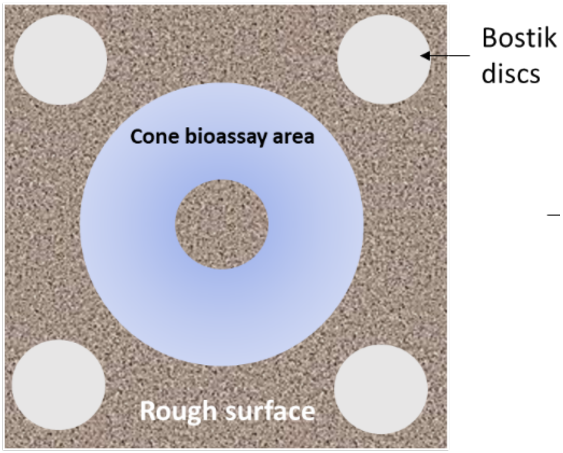


**Figure 1. Method of sampling insecticide-treated surfaces with wettable powder formulations.**

1. Determine the area to be tested as indicated in Figure 1.
2. Four Bostik per sample should be placed next to the cone bioassay test area, as shown in Figure 1.
3. Ensure good contact (finger rubbing) for approximately one minute (using gloves, do not touch surface).
4. Pull off the Bostik discs and combine with filter paper to prevent sticky material from self-folding or overlapping during storage and shipping.
5. In each filter, record the sample number with a pencil, but do not write in the disc area.
6. Wrap disc samples in tin foil and indicate sample number using permanent marker on the tin foil.
7. Provide a sample login sheet in excel that contains detailed information about each sample sorted by sample number, and sampling date. Please note that data will be reported on the same excel file once the analysis is completed.
8. Samples should be stored in the dark at 4$^{o}$C until they are shipped to LSTM.

### Wall cone bioassays:

- Place mosquitoes in batches of 10 into paper cups prior to the home visits.
- Cone bioassays on wall surfaces will be conducted on wall surfaces such as plastered and painted, plain brick and mud-walled using a standardized WHO protocol (WHO 2006).
- The study will use wall surfaces available in the 60 cohort houses with sampling conducted every 2 months for a period of up to 12 months.
- Following standard WHO methodology, cones will be placed at heights of 1.0 m above the floor.

*Sampling protocol*

- Cones lined with self-adhesive tape will be fixed on the sprayed walls for the assay. The control cone will be affixed to a wall lined with a paperboard.
- Five to 10-day-old female mosquitoes will be used for the tests. Mosquitoes both susceptible and wild *An. gambiae s.s.* will be introduced into the plastic cones in batches of 10 and left exposed on the sprayed surface for 30 minutes.
- Record knockdown resistance after 1hr and then keep mosquitoes for 24h to 48 hrs and record mortality.
- Outcome: This test will inform us whether there is sufficient insecticide on the wall to kill lab susceptible mosquitoes. This is a quality assurance assay.
- Store dead mosquitoes in silica gel and alive mosquitoes in RNALater and label.
