## Supplementary material for "Significant variations in tolerance to clothianidin and pirimiphos-methyl in *Anopheles gambiae* and *Anopheles funestus* populations during a dramatic malaria resurgence despite sustained indoor residual spraying in Uganda": Protocol for sampling walls for Actellic

*Sampling protocol*

**<u>Sampling protocol of Households sprayed with Actellic® 300CS insecticide:</u>**

1. There are 60 houses to be monitored with the cohorts.

2. We plan to sample insecticide wall content every 3 months giving us a total of 4 sampling rounds. We also plan to sample from two separate rooms (preferably, sitting room and bedroom) within the same household.

3. Sampling will be done at time 0 (during/shortly after spray), time 3 (three months after spraying). Time 6 (6 months after spraying), time 9 (9 months after spraying) and time 12 (12 months after spraying).

4. Samples will be sent to LSTM immediately after each round for analysis.

5. The sampling on the walls will be done as follows:

a) At time 0, three filter papers will be placed in the upper (U), middle (M) and Lower (L) part of each room prior to the spray. To ensure accurate data collection, a total of 6 filter papers will be provided per household.

b) After the spraying process is completed, the surface is allowed to dry, which typically takes about 2-3 hours. Once the surface is completely dry, the filter papers will be removed from the sprayed surface. This step is essential as it ensures that the filter papers do not stick to the surface and can be easily peeled off without causing any damage or leaving any residue behind. Therefore, it is important to follow the recommended drying time before removing the filter papers.

c) The side of the filter paper that was exposed to the spray nozzles should be covered with Sellotape, wrapped in tin foil, and labelled with the corresponding sample number.

d) After labelling the filter papers, it is important to ensure that the corresponding sample number is correctly recorded in the samples log sheet. This will help avoid confusion and errors during the analysis phase of the experiment. To make the process more efficient, it is recommended to use a standardized template, such as the one shown in table 1, to populate the samples log sheet with all relevant information, such as the date, time, and location of the sample collection.

Table 1 Sample info

| Sample name | Spray Date | Date sample taken | Spray operator ID | House ID | Type of surface | Location | Sample | Concentration (g/m2) | Mean [P-Methyl] g/m2 per Room | SD | Mean [P-Methyl] g/m2 per House | SD | LSTM Code |
|---|---|---|---|---|---|---|---|---|---|---|---|---|---|
| 1 | 25/05/2019 | 31/05/2019 | BO 196 | M0337S035E001P01 | Wood | Living room | Upper | | | | | | 1 |
| | | | | | | | Middle | | | | | | 2 |
| | | | | | | | Bottom | | | | | | 3 |
| | | | | | | Bedroom | Upper | | | | | | 4 |
| | | | | | | | Middle | | | | | | 5 |
| | | | | | | | Bottom | | | | | | 6 |

e) This labelling process is critical for accurate data analysis and we kindly request that all participants follow this procedure.

f) The area that contained the filer papers should be marked by pencil or chalk.

g) After collecting the filter paper samples, the next step is to carefully position four adhesive dots in close proximity to each filter paper position. The purpose of this is to effectively extract and analyze the insecticide wall content. An example of how this is done is shown in Figure 1. By using this methodology, a total of six samples will be obtained for each household. This will include four glue dots per sample. It is important to ensure that the adhesive dots are positioned accurately and precisely to obtain accurate results during analysis.

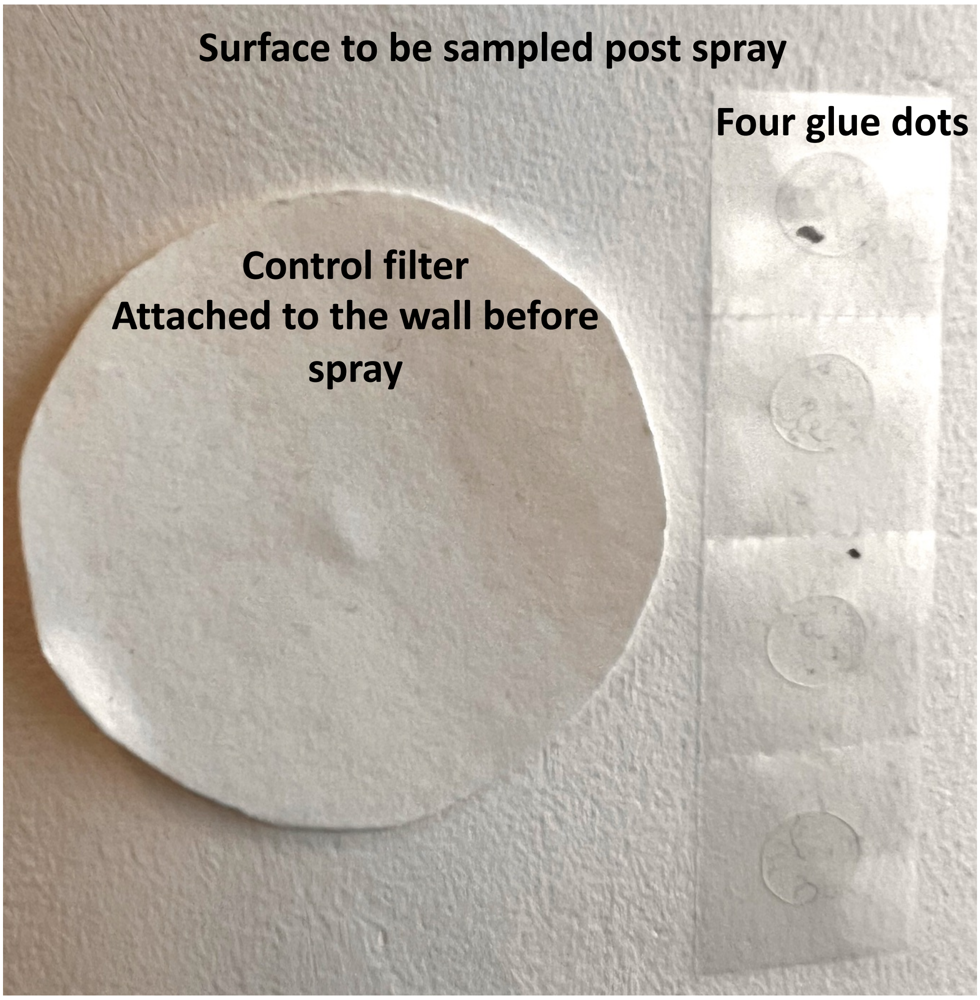


Figure 1: Sampling white painted wall with four glue dots post spray with Actellic 300SC

h) To ensure proper identification of the samples, four glue dots will be carefully placed in the filter paper and labeled accordingly. These labels will include U, M, L, and the room where they were collected, as depicted in Figure 2.

*Sampling protocol*

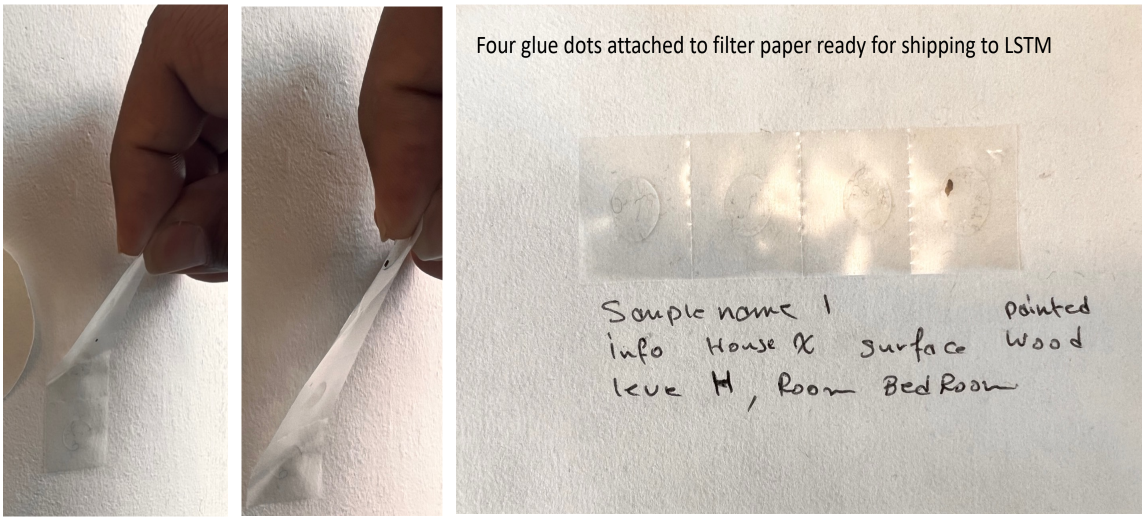


Figure 2: Glue dots are removed and attached to filter paper labelled with sample info

i) At the designated time period between 3 and 12, an additional rounds of glue dots sampling will be performed, following the procedure described in steps 5g to 5h.

6. During the sampling on the walls:

a) Determine the area to be sampled.
b) Four glue dot per part of the wall should be placed next to the area that has a cone and control filter paper.
c) Ensure good contact (finger rubbing) for approximately one minute (using gloves, do not touch surface).
d) Pull off the glue dots carefully to make sure that glue dots still attached to glossy back and combine with filter paper to prevent sticky material from self-folding or overlapping during storage and shipping.
e) In each filter, record; Batch; Spray Date; Date sample was taken; House ID; Type of surface; Location; Sample (part of wall) as indicated in table 1.
f) Wrap samples (control filter and glue dots) in tin foil and indicate sample number using permanent marker on the tin foil.
g) Provide a sample login sheet in excel that contains detailed information about each sample sorted by sample number, and sampling date.
h) Please note that data will be reported on the same excel file once the analysis is completed.
i) Samples should be stored in the dark at 4°C and shipped to LSTM.

7. Wall cone bioassays:

a) Place mosquitoes in batches of 10 into paper cups prior to the home visits.
b) Cone bioassays on wall surfaces will be conducted on wall surfaces such as plastered and painted, plain brick and mud-walled using a standardized WHO protocol (WHO 2006).
c) The study will use wall surfaces available in the 60 cohort houses with sampling conducted every 2 months for a period of up to 12 months.
d) Following standard WHO methodology, cones will be placed at heights of 1.0 m above the floor.
